# Pattern formation and bistability in a synthetic intercellular genetic toggle

**DOI:** 10.1101/2022.08.02.502488

**Authors:** Bárbara de Freitas Magalhães, Gaoyang Fan, Eduardo Sontag, Krešimir Josić, Matthew R. Bennett

## Abstract

Differentiation within multicellular organisms is a complex process that helps to establish spatial patterning and tissue formation within the body. Often, the differentiation of cells is governed by morphogens and intercellular signaling molecules that guide the fate of each cell, frequently using toggle-like regulatory components. Synthetic biologists have long sought to recapitulate patterned differentiation with engineered cellular communities and various methods for differentiating bacteria have been invented. Here, we couple a synthetic co-repressive toggle switch with intercellular signaling pathways to create a “quorum-sensing toggle.” We show that this circuit not only exhibits population-wide bistability in a well-mixed liquid environment, but also generates patterns of differentiation in colonies grown on agar containing an externally supplied morphogen. If coupled to other metabolic processes, circuits such as the one described here would allow for the engineering of spatially patterned, differentiated bacteria for use in biomaterials and bioelectronics.

## Introduction

One goal of synthetic biology is the creation of engineered biological systems that can perform a wide variety of functions (Benner & Sismour, 2005; Cameron et al., 2014; Cheng & Lu, 2012; Church et al., 2014). Such systems could be used for various environmental, industrial, and medical applications (Callura et al., 2012; Khalil & Collins, 2010; McCarty & Ledesma-Amaro, 2019; Xia et al., 2019). However, synthetic biology is also advancing basic research by providing a bottom-up approach to understanding phenomena governed by nontrivial genetic regulatory mechanisms. This is done by creating and perturbing “synthetic gene circuits” in living systems that behave similarly to their natural counterparts and can thus serve as a model systems (Davies & Glykofrydis, 2020; Weisenberger & Deans, 2018).

As part of the ground up approach to basic research, synthetic biologists have long sought to use engineered cells to recapitulate spatial patterns seen in multicellular systems (Cachat et al., 2017; Davies & Glykofrydis, 2020; Grant et al., 2020; Kim et al., 2020; Santos-Moreno & Schaerli, 2018; Sekine et al., 2018). Many mechanisms have been proposed to explain the appearance of natural patterning, such as the reaction-diffusion model (Turing, 1952) and the positional information model (or French flag model) (Wolpert, 1969). These mechanisms have been used as blueprints for synthetic analogs of biological patterns (Diambra et al., 2015; Karig et al., 2018a; Sekine et al., 2018). Scientists are also working towards self-organizing patterns (Cachat et al., 2017; Cao et al., 2016; Curatolo et al., 2020; Liu et al., 2011; Payne et al., 2013; Potvin-Trottier et al., 2016; Santos-Moreno & Schaerli, 2018) that use intercellular signals to regulate transcriptional activity (Cao et al., 2016; Curatolo et al., 2020; Liu et al., 2011; Payne et al., 2013).

Indeed, the circuit topology of the first synthetic gene circuit to be described, the “genetic toggle switch” (Gardner et al., 2000a), has also been implicated in spatial patterning. The toggle switch is comprised of just two repressors that regulate each other’s promoters – and it therefore has two possible transcriptional states corresponding to one active repressor gene, and the other repressed. Gene networks akin to the toggle switch are thought to help stabilize and refine spatial boundaries within differentiated populations because the two expression states of the toggle are generally mutually exclusive (Briscoe & Small, 2015). Toggle-like regulatory components are found throughout developmental processes, such as in the anterior-posterior development of the *Drosophila* blastoderm (Nasiadka et al., 2002; Struhl, 1989), and in the dorsal-ventral development of the vertebrate neural tube (Alaynick et al., 2011; Dessaud et al., 2008).

In multicellular systems, toggle switches can function independently. In the absence of any external signal (e.g., morphogens), the internal stochastic dynamics and bias of each cell determine the transcriptional states of the respective toggle switches. In other words, each cell randomly assumes one of the two states. However, in the presence of an external morphogen gradient, a distinct boundary is formed between cells in the two different transcriptional states. This can occur if the toggle switch exhibits hysteresis (bistability) as a function of the morphogen.

Here, we built a version of the genetic toggle switch in *Escherichia coli* that uses intercellular signaling to reinforce each transcriptional state. For instance, if a cell is in the “ON” state it produces an intercellular signal that up-regulates the ON state in nearby cells. The addition of intercellular signaling to the toggle has been proposed as a means of creating a population-level toggle switch, allowing all cells in the population to simultaneously reside in one of the two possible transcriptional states (Nikolaev & Sontag, 2016). We experimentally confirm that, given the right conditions, this version of the toggle does exhibit population-level bistability in a well-mixed liquid culture. Additionally, we show that when grown on solid agar imbued with an exogenous morphogen, colonies of these cells form three dimensional patterns that are distinctly different from those observed in colonies of cells containing a traditional toggle switch lacking intercellular signaling. In particular, the addition of intercellular signaling creates a regime that enables a shift in the resulting pattern. We develop a mechanistic mathematical model of the system, to explain how degradation, diffusion, and sequestration of the signaling molecules and inducers determine the observed patterns.

## Results

### General characteristics of QS and NQS toggles

Here, we call the version of the toggle switch that includes intercellular signaling the “QS toggle”, as it uses refactored **q**uorum **s**ensing pathways to generate intercellular signals. The QS toggle is a version of the genetic toggle switch (Gardner et al., 2000a) that includes two repressors that repress each other’s promoters (LacI and TetR). Additionally, it includes a reporter for each state (YFP and CFP). The circuit also includes two orthogonal QS pathways (CinR/I and RhlR/I) (Chen et al., 2015; Lithgow et al., 2000; Pesci et al., 1997) to produce the necessary intercellular signaling. A representation of the QS toggle’s two fluorescent states is shown in Fig. 1A. For the yellow state to be active (Fig. 1A, left), LacI represses the promoters driving the expression of *tetR, cfp* and *rhlI*. With *tetR* repressed, *cinI* is expressed, and its protein catalyzes the production of C14-HSL. When bound to the CinR transcription factor, C14-HSL activates the expression of *lacI* and *yfp*. Alternatively, for the blue state to be active (Fig. 1A, right), TetR represses the promoters driving the expression of *lacI, yfp* and *cinI*. With *lacI* repressed, *rhlI* is expressed, and its protein produces C4-HSL. When bound to the RhlR transcription factor, C4-HSL activates the expression of *tetR* and *cfp*. One can exogenously induce the yellow state by adding anhydrotetracycline (aTc), which will inactivate TetR (Fig. 1E). Similarly, one can exogenously induce the blue state by adding isopropyl-β-D-1-thiogalactopyranoside (IPTG), inactivating LacI (Fig. 1D). The QS toggle is unbalanced in the absence of inducers: it exhibited a preference to the blue state (Fig. 1D, E). We also found that this imbalance is dependent on the QS network (Fig. S2I, J). The interaction of the main circuit elements is summarized in Fig. 1C. The QS toggle topology (Fig. 1C, left) differs from the conventional (**n**on-**q**uorum **s**ensing toggle or “NQS toggle”-Fig. 1C, right) topology by the presence of two QS networks, creating extra positive feedback loops. The NQS circuit includes the same repressors and reporter genes but driven by promoters that are only responsive to these repressors (Fig. 1B). The NQS toggle can also be tuned with IPTG and aTc (Fig. 1F and 1G). For all experiments, we transformed the plasmid-borne QS and NQS circuits into *E. coli* cells that contained constitutively expressed *cinR* and *rhlR* in their genome.

**Figure 1:**
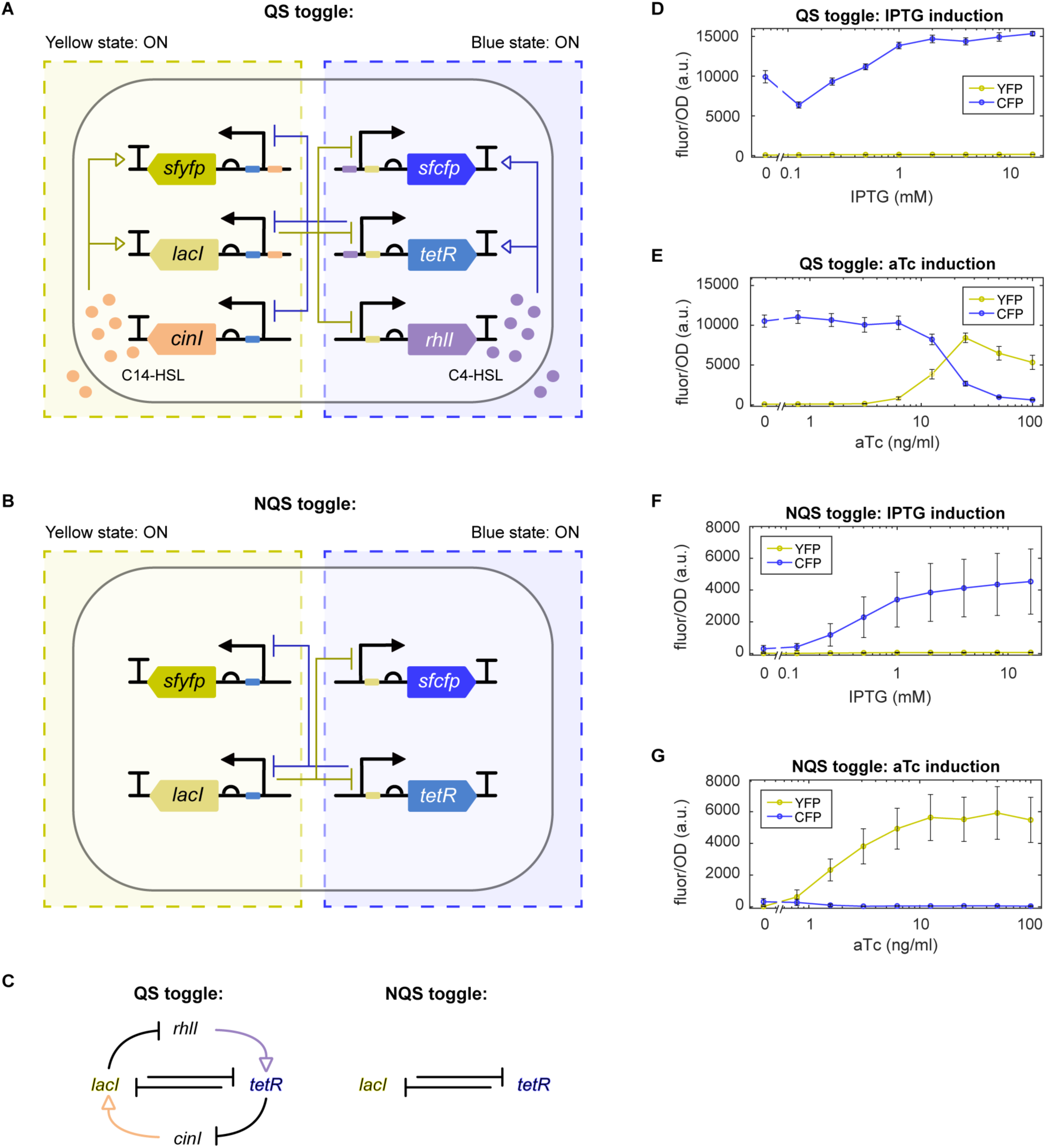
QS and NQS toggle circuit design. For both circuits, IPTG can induce the blue state, while aTc can induce the yellow. **A)** QS toggle: When the yellow state is ON, LacI represses *tetR, rhlI* and *sfcfp* genes, while C14-HSL activates *lacI* and *sfyfp* genes. When the blue state is ON, TetR represses *lacI, cinI* and *sfyfp* genes, while C4-HSL activates *tetR* and *sfcfp* genes. **B)** NQS toggle: When the yellow state is ON, LacI represses *tetR*, and *sfcfp* genes. When the blue state is ON, TetR represses *lacI* and *sfyfp* genes. **C)** The QS and NQS toggle topologies differ by the presence of 2 QS networks. **D)** QS toggle induction with IPTG in liquid culture. **E)** QS toggle induction with aTc in liquid culture. **F)** NQS toggle induction with IPTG in liquid culture. **G)** NQS toggle induction with aTc in liquid culture. Lines represent the average fluorescence and error bars represent the standard deviation of 3 technical replicates for 3 independent experiments.

We first checked whether cells containing the QS toggle exhibited population-level bistability. Genetic toggles are often bistable (Barbier et al., 2020; Gardner et al., 2000a; Lugagne et al., 2017; Wang et al., 2009), *i*.*e*. they can stably reside in either of two possible transcriptional states. To find the bistable region for both the QS and NQS toggles, we grew overnight cultures with either IPTG or aTc to allow for the populations to start in either state (blue or yellow, respectively). We then grew these cells in liquid culture using various combinations of IPTG and aTc concentrations and used flow cytometry to measure the CFP and YFP fluorescence of individual cells. After three hours, we observed that, when starting in the yellow state (*i*.*e*. previously grown in media with aTc), QS toggle cells all remained in the yellow state for all combinations of inducer (Fig. 2A, right). However, when starting in the blue state (i.e., previously grown in media with IPTG), QS toggle populations shifted to the yellow state at sufficiently high aTc but remained in the blue state at lower concentrations (Fig. 2A, left). Importantly, we observed cultures in which all cells were either in the blue or yellow state depending on the starting condition and the inducer concentration (Fig. 2A) – indicating that the QS toggle is bistable at the population level in those conditions. Meanwhile, NQS toggle cells transitioned to blue and yellow, when starting from the opposite color initial conditions (Fig. 2B). We also observed that cells pre-induced with IPTG primarily did not show any fluorescence (Fig. 2B, left), although the blue state was inducible. A third state (OFF), in which cells exhibited little to no fluorescence of either type, was more common within NQS cells. The NQS toggle was also bistable for some conditions. When growing the cells for 9 hours instead, the NQS toggle showed a decrease in both intensities in all tested conditions, while the QS toggle tended to show stronger fluorescence intensities (Fig. S1).

**Figure 2:**
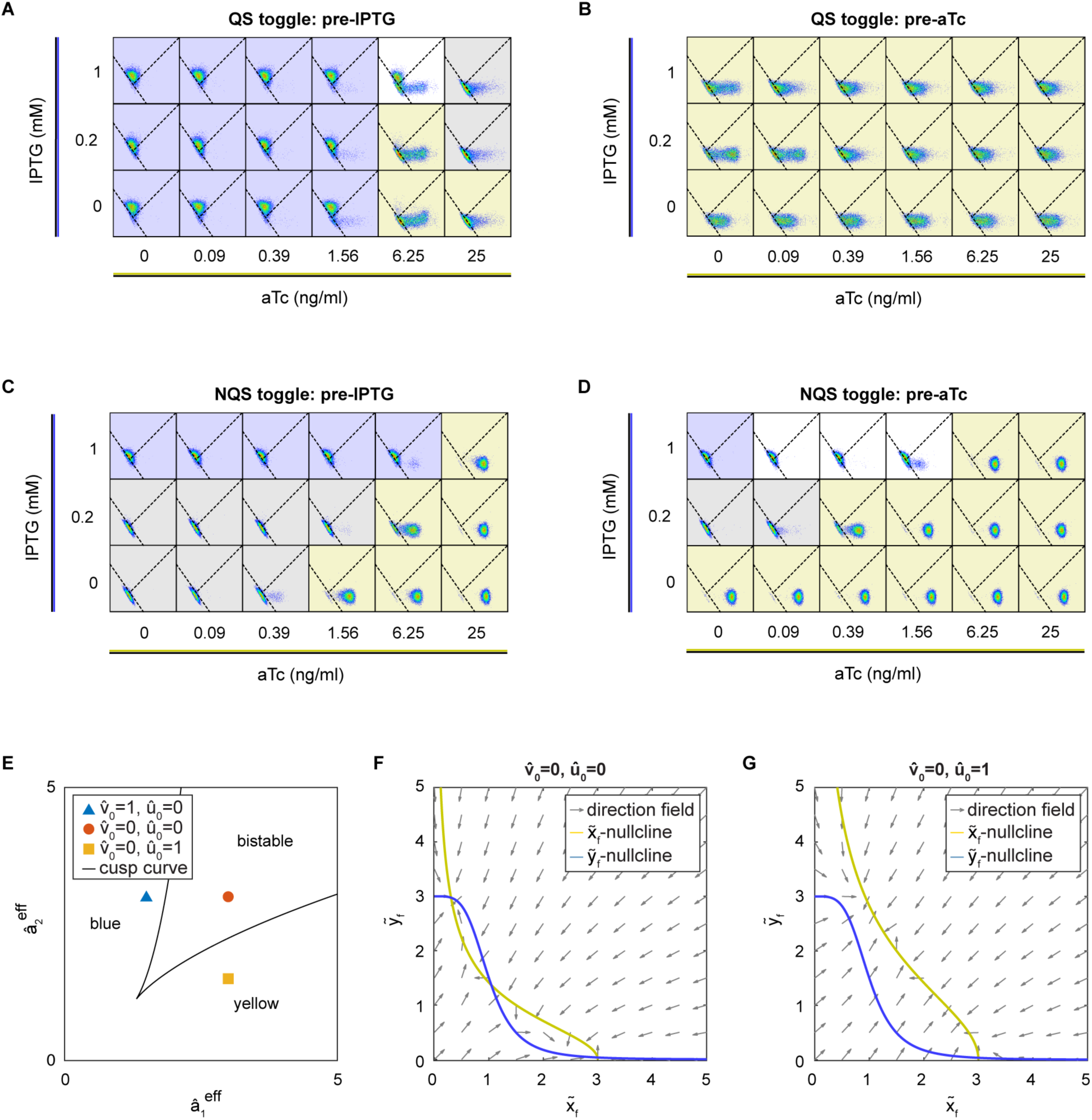
Behavior of QS and NQS toggle cells when treated with a combination of inducers. **A-B)** Flow cytometry data of QS toggle cells that were pre-induced with either IPTG (A) or aTc (B). Each dot is a single cell classified within a gate. Gates were determined with single color and double negative controls. Dashed lines in each plot represent the boundaries between the three distinct gates, which represent cellular states: CFP+ (top gate), YFP+ (bottom-right gate), and OFF (bottom-left gate). Background colors in each plot represent which state the majority of cells are in (>50%): blue color indicates mostly CFP+ cells, yellow plots are mostly YFP+, gray plots are mostly OFF, and white plots indicate cells that are present in multiple states (<50% each). **C, D)** Flow cytometry data of NQS toggle cells that were pre-induced with either IPTG (C) or aTc (D). **E)** Bifurcation diagram of the QS toggle system over the nondimensionalized parameters 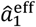 and 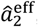, the effective promoter strength of the repressors. Here we demonstrate a case where without exogenous inducers, aTc or IPTG, the system is bistable (red dot). Adding aTc or IPTG (blue triangle and yellow square respectively), lowers the effective promoter strength, which can in turn lead to changes in the state of the cell to be monostable in either the blue or yellow state. **F, G)** Phase portrait of the QS toggle system for chosen parameter values denoted by the red dot and blue triangle in E. Variable and parameters in E-G are nondimensionalized (see STAR methods).

We next developed a mathematical model of the toggle switches to understand under what conditions the two versions of the circuit can exhibit bistability. Denoting by x(t) and y(t) the concentration of LacI and TetR, respectively, and by *g*(*t*) and *h*(*t*) the intracellular concentration of QS signals, C14-HSL and C4-HSL, the model takes the form

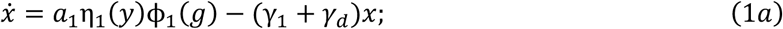

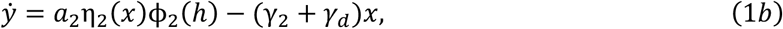

where the parameter *a*_*i*_ determines the maximal production rates, *γ*_*i*_ the individual degradation rates for LacI and TetR (numbered *i* = 1,2, respectively), and *γ*_*d*_ the rate of dilution due to cell growth. We model repression and activation of the promoters using Hill functions, *η*_i_ and ϕ_i_, respectively. The intracellular concentration of QS signals obeys similar equations (See STAR methods for the full model).

We nondimensionalized Eq. (1) and performed a bifurcation analysis (see STAR methods), to first confirm the existence of the region of bistability in the QS toggle system model (See Fig. 2E). Moreover, as is evident in the production term of LacI and TetR in Eq. (1), the preferred state of the QS toggle is not determined only by the presence of the QS signals, but also the relative promoter strength. This is characterized by the following nondimensionalized variables,

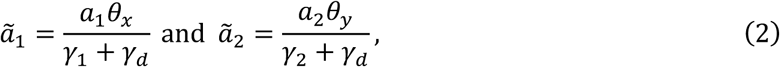

where *θ*_*x*_ and *θ*_*y*_ are the repression thresholds for LacI and TetR, respectively, used in the Hill functions, *η*_i_. When exogenous inducers, aTc or IPTG, are added to the system, the relative promoter strength is modified. Following experiments, we assumed that these inducers are provided at some constant level, denoted by nondimensionalized parameter 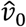 and 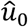. This leads to the following effective promoter strengths,

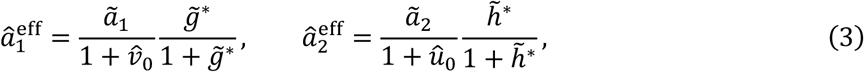

with 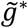 and 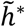 representing the nondimensionalized C14 and C4 concentrations in the system. Depending on the values of the bifurcation parameters 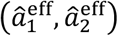 the system can be either mono-or bistable (see Fig. 2E-G; STAR methods for the full model).

The mathematical model thus predicted that bistability depends on the QS network and relative promoter strengths. We tested this prediction experimentally by engineering a few variants of the QS toggle that changed those factors (see SI). These variants exhibited changes in state preference, agreeing with the prediction of the model (Fig. S2).

### QS and NQS toggle behavior in colonies

We next asked how the QS toggle behaves in an environment in which the QS signals are not homogeneously distributed in the population, allowing for spatial patterns to arise. To do this, we grew QS and NQS toggle colonies in LB agar plates containing aTc and monitored their growth and fluorescence over time.

Most strikingly, we noticed that many QS toggle colonies formed a blue ring surrounding a yellow disc (Fig. 3A), with the size of the yellow disc correlated with the aTc concentration in the agar (Fig. 3B). ATc is known to be temperature sensitive (Politi et al., 2014). Since aTc induces the expression of the yellow state, we hypothesized that aTc degradation was driving the emergence of blue cells. Indeed, we verified that prior incubation of aTc plates at 37°C directly alters the blue ring size and time of appearance (Fig. S3). Thus, the decay of aTc creates a temporal morphogen gradient, resulting in a switch in the state of some of the population.

**Figure 3:**
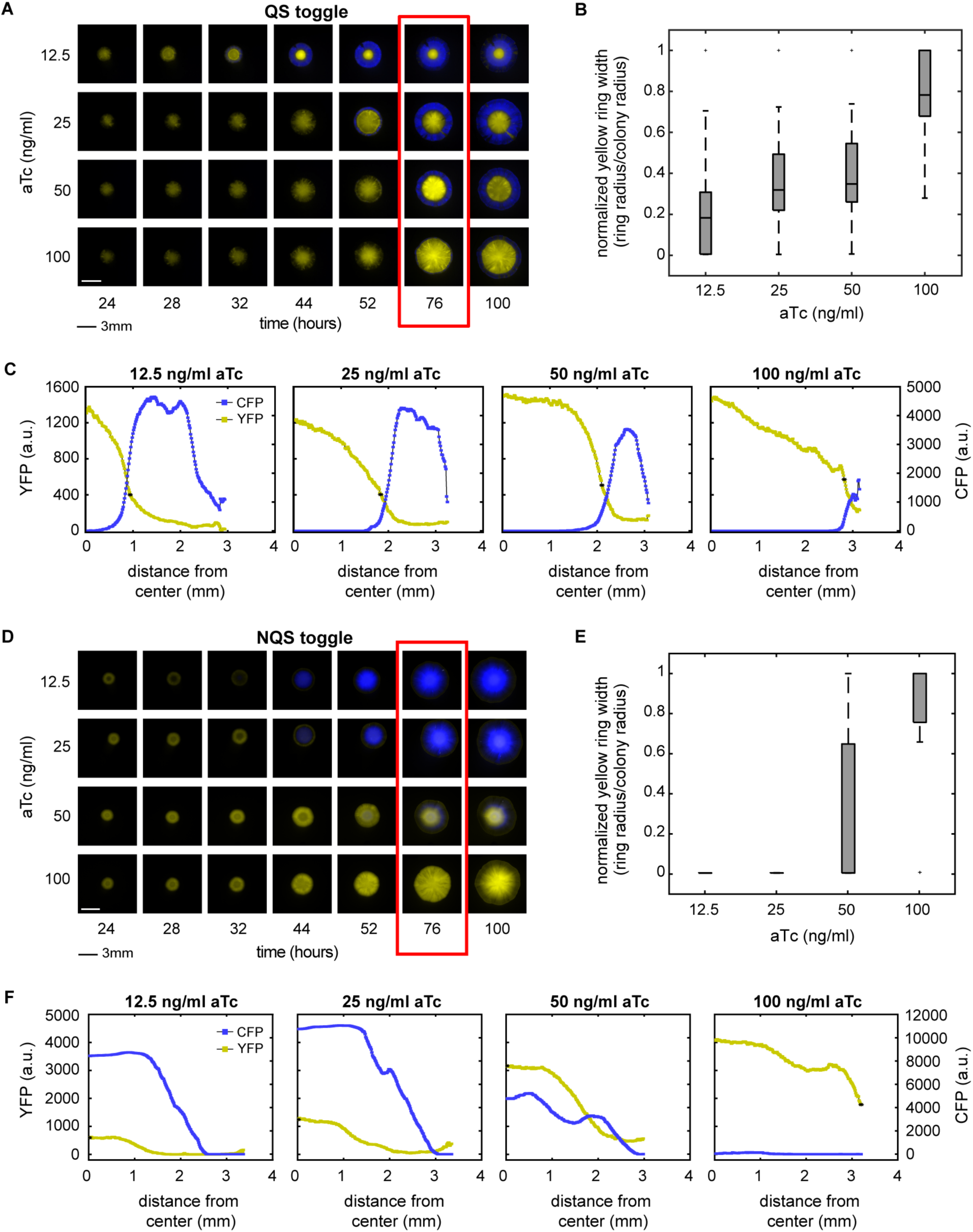
Expanding QS and NQS toggle colonies show aTc-dependent patterning. **A)** QS toggle colonies obtained from plates with different initial aTc concentrations, shown at different times. **B)** Yellow ring width (center disc) was measured among all colonies to quantify the pattern. The width was determined as the distance from the center to the threshold point, at which yellow fluorescence fell below 30% of its maximum. The only pairwise combination of widths not found to be statistically different was 25 and 50 ng/ml aTc (*p* < 0.01, Kruskal-Wallis non-parametric test). Data represents at least 11 independent experiments. **C)** Fluorescence intensity cross-sectionals of the colonies shown in red in (A). Curves represent the average fluorescence of 4 radii (90° apart) of the same colony. **D)** NQS toggle colonies obtained from plates with different initial aTc concentrations, shown at different times. **E)** Yellow ring width was used to quantify the patterns as in (B). We considered yellow ring width to be zero when blue cells dominate the center (see methods for full set of assumptions). All pairwise that included 100 ng/ml aTc were found to be statistically different (\**p* < 0.01, Kruskal-Wallis non-parametric test). Data represents 2 independent experiments. **F)** Fluorescence intensity cross-sectionals of colonies shown in red in (C). Curves represent the average fluorescence of 4 radii of the same colony.

In the images of QS toggle colonies, we observed little overlap between cells in the yellow and blue states (Fig. 3C). Over time, the blue ring generally expanded as the colony grew, but the boundary between the yellow and blue cells remained roughly fixed. Wherever cells in the blue state took over, yellow fluorescence tended to decrease.

When grown on solid media, the NQS toggle behaved differently from the QS toggle. Like the QS toggle, NQS toggle colonies were initially yellow when the media contained a high enough concentration of aTc. Cells in the blue state emerged eventually but did so in a way distinct from what we observed in the QS toggle: Blue cells first appeared in colony centers, and did so earlier and at lower aTc concentrations (Fig. 3D). Most NQS toggle colonies grown with 12.5-50 ng/ml aTc showed blue cells in the center at 76h (Fig. 3E), which resulted in yellow region width measurements close to zero (see Methods).

One of the main differences in spatial patterns between the two toggles was the degree of radial separation between the states. The QS toggle colonies appeared to segregate well in contrast to the NQS toggle colonies (Fig. 3F). To quantify the level of segregation, we measured the pixel color overlap in images from both QS and NQS colonies: For each pixel, we measured the intensity of both blue and yellow fluorescence and in Fig. 4A and B (bottom panels), we show the normalized YFP and CFP fluorescence as a function of distance from the center of the colony. We said that colors overlapped in a pixel when the values of both YFP and CFP fluorescence intensities were each above a threshold of 0.3 (out of a maximum intensity normalized to 1). To obtain the relative overlap count for a colony, we divided the number of overlapping pixels by the total number of pixels covering the colony radius (See Fig. 4C).

**Figure 4:**
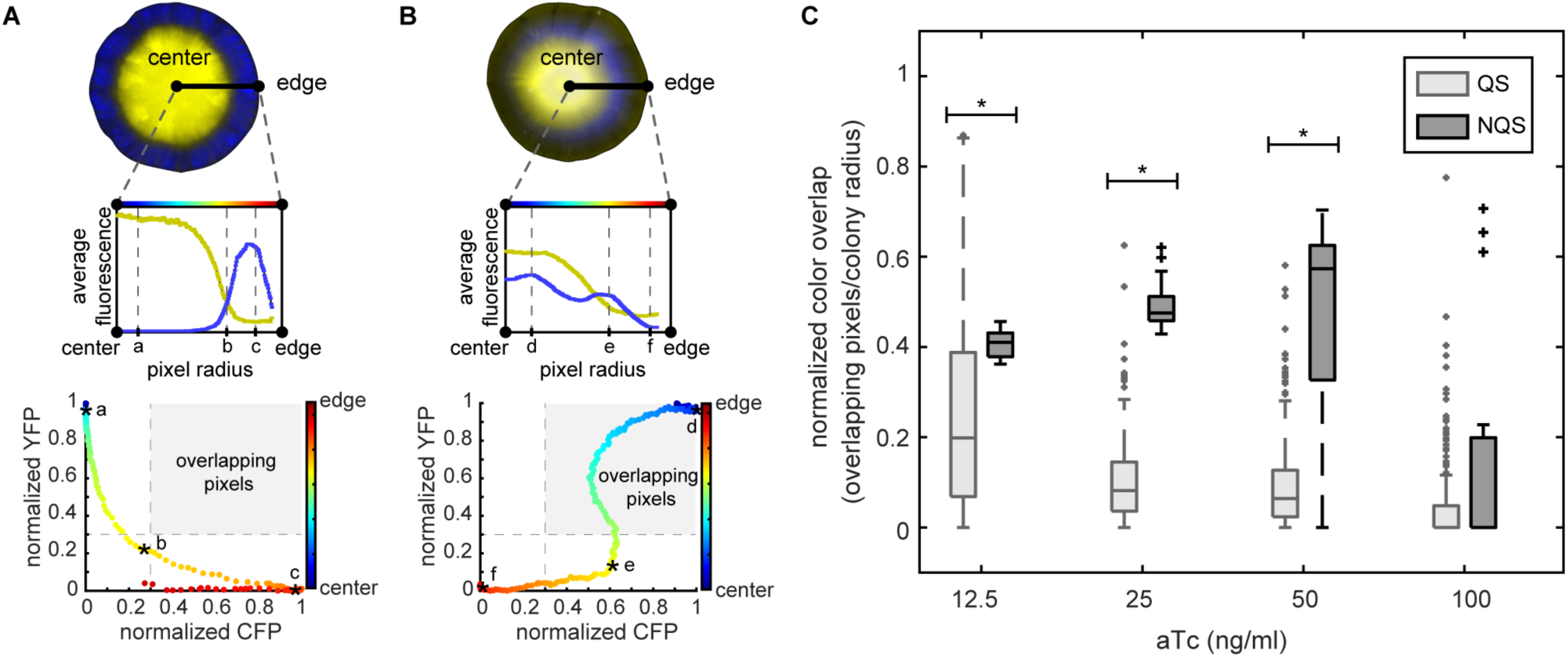
QS colonies create better radially separated patterns than NQS colonies. Measurement of color overlap was used to quantify the spatial segregation of states in each colony. Average fluorescence of each colony’s radii plotted from the center of the colony to the edge (A and B, top plots). Each pixel was also plotted for both normalized fluorescence values (A and B, bottom dot plots). Pixels were classified as overlapping when both normalized fluorescence values were above a predefined threshold (inside the gray boxed region). **A)** Example of the typical behavior of QS colonies. The QS colony shown is the same as that in Fig. 3A, at 50 ng/ml and 76 hours. Asterisks (a, b, c) show the same pixels in both plots. **B)** Example for the typical behavior in NQS colonies. NQS colony shown is from Fig. 3C, 50 ng/ml at 76 hours. Asterisks (d, e, f) show the same pixels in both plots. **C)** Quantification of pixel overlap for QS (light gray) and NQS (dark gray) colonies. The number of overlapping pixels was normalized by the colony radius (total). \**p* < 0.01, Mann-Whitney non-parametric test. Data for each QS test contains over 180 colonies from at least 11 independent experiments, while data for each NQS test contains at least 15 colonies from 2 independent experiments.

Color overlap was significantly higher in the NQS compared to QS colonies in 12.5, 25, and 50 ng/ml aTc plates (*p <* 0.01, Mann-Whitney non-parametric test). In 100 ng/ml aTc plates, we observed little overlap with both circuits, mostly because not all colonies contained cells in the blue state. When we excluded from analysis colonies with cells in only one state, NQS colonies displayed higher overlap than QS colonies at all aTc concentrations (Fig. S4).

Lower overlap indicates a better separation of colors, and hence a better spatial separation of cells in the two states along the radial direction of a colony. Therefore, QS cells in different states were better segregated radially than NQS cells. However, since we imaged the colony from above, the higher degree of overlap in the NQS case could indicate either that cells in the two states are intermingled, or that they are segregated vertically. To test which of these hypotheses is more likely, we extended our mathematical models to include spatial effects, such as colony growth, and the diffusion of inducers and signaling molecules within the agar and the colony. We assumed that the colony is conical and grows through the addition of cells in an active growing zone at the interface of the colony with agar (Warren et al., 2019) (see STAR methods).

Simulations of our expanded models recapitulated experimental observations, and suggested answers to our questions about the observed segregation of cells in different states. In the simulations of the NQS toggle colony (Movie S1), with smaller colony sizes and abundant aTc supply from the agar, aTc levels are relatively high everywhere in the NQS toggle colony. This explains the initial colony-wide yellow state we observed in experiments (Fig. 3D). In this initial yellow state, intracellular concentration of TetR is low, leading to a low sequestration rate of aTc inside the colony. The top of the colony is further from the aTc source (agar), so that a top-down aTc gradient forms over time. As the model colony grows, we observe an increase in sequestration and degradation of aTc (due to 37°C incubation over time). Eventually, the aTc level at the top of the colony, where concentration is lowest, degrades below the point needed to keep cells in the yellow state, causing the top of the colony to turn blue. Cells entering the blue state produce more TetR, which enables higher aTc sequestration, further lowering surrounding aTc levels. As a result, we observed a blue wave traveling downwards from the top of the colony. Thus, our model predicted that the two states are vertically segregated in NQS colonies, suggesting that spatial segregation, rather than heterogeneity, is responsible for the experimentally observed color overlap. We also examined the state of the system by tracking the bifurcation parameters (Eq. 3) at the center (top) and the periphery (rim) of the NQS colony over time (Fig. 5D). In simulations, the top of the colony moves from the bistable state into the blue state, while the rim of the colony remains in the yellow state (Fig. 5B, C). Our simulations and bifurcation analysis also revealed that at higher initial aTc concentrations the blue state takes over from the top at a later time (Fig. 5A), as it takes longer for aTc to degrade to the hysteresis point at which cells can switch states.

**Figure 5:**
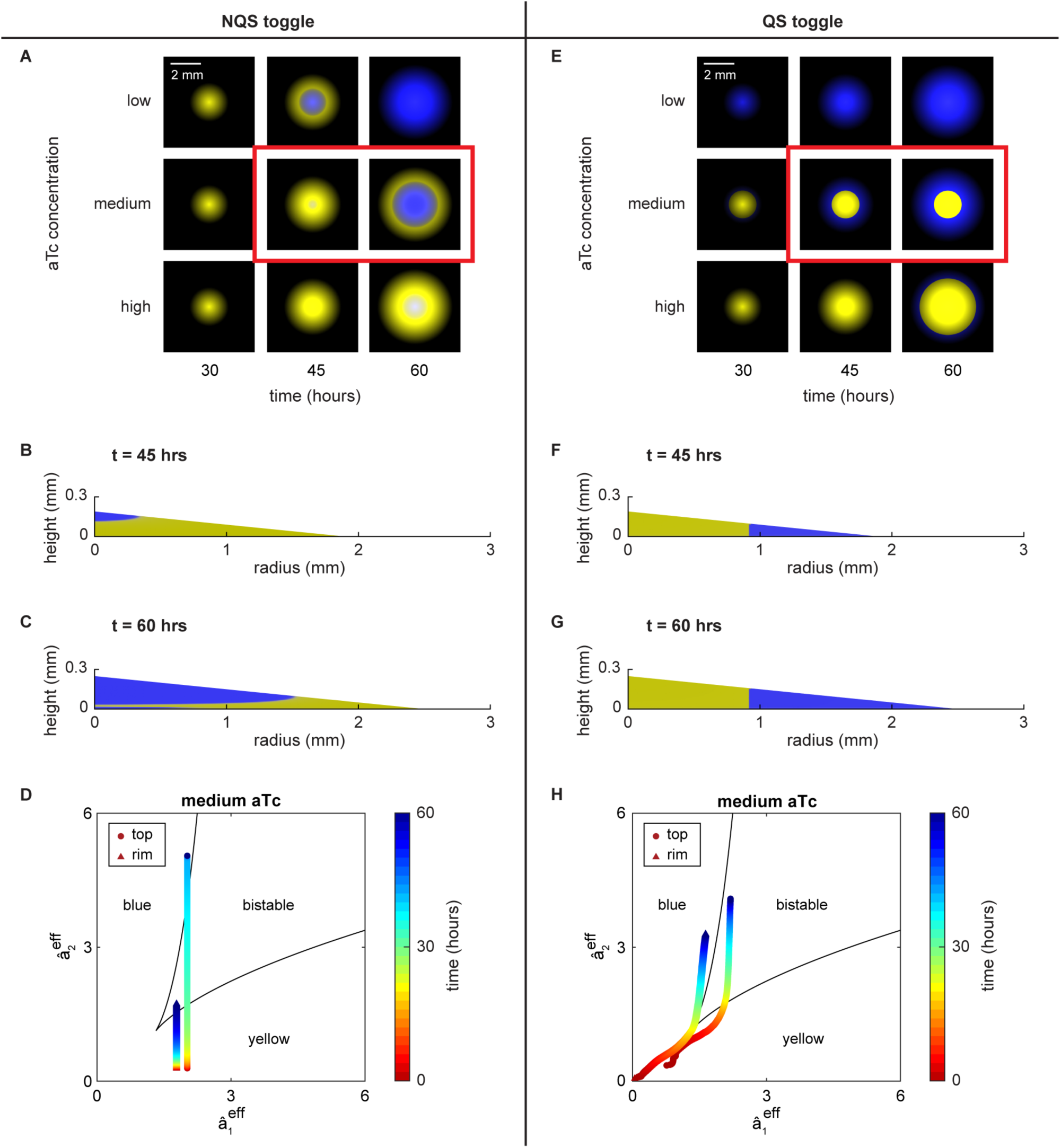
PDE simulations of NQS and QS toggles. **A)** Simulations of the NQS colony at different times and initial aTc concentration show the emergence of cells in the blue cells at the center of the colony. **B)** The cross-sectional view of the model colonies shows that cells enter the blue state first at the top of the colony, and that cells in the two states remain vertically segregated. **C)** Bifurcation diagram showing the bifurcation parameters’ trajectories of cells at the top (circle) and rim (triangle) of the NQS colony. The top of the colony flips from yellow to blue. **D)** Simulations of a model QS colony show that a blue ring emerges dependent on the initial aTc concentration. **E)** The cross-sectional view of the model colonies shows that the cells in the two states are radially segregated, and that the boundary between the two states is roughly fixed, in agreement with experiments. **F)** Same as in C), but showing that in the QS colony the rim of the colony flips from the yellow to blue state.

In simulated QS toggle colonies (Movie S2), when the colony is small, abundant aTc and basal C14-HSL production led to the establishment of the initial yellow state across the colony. This state was then reinforced by the high C14-HSL signal. As indicated by Eq. 2, the dilution rate alters the effective promoter strength, leading to different steady states. Cells that are not growing or growing slowly, such as those at the top of the colony, stay in the bistable region, and do not switch states (Fig. 5H). This explains why the center of the colony remains yellow, despite reduced aTc levels due to degradation and sequestration. In contrast, for fast-growing cells, the drop in aTc concentration and the slow accumulation of C4-HSL signal from basal production allow cells to enter the blue state (Fig. 5F). Cells in the blue state start to appear at the periphery of the colony, where cells grow fastest (Movie S2). This explains why in the QS toggle color overlap is small (vertical slices are in the same state), and the boundary between the yellow and blue states is maintained at the point where the first cells turn blue (Fig. 5F, G), in agreement with experimental observations (Fig. 3A). Like the NQS toggle, higher aTc concentrations lead to the blue ring emerging later in time (Fig. 5E).

To verify the predictions of the model about the different spatial structures in QS and NQS colonies, we used confocal microscopy to image the 3D structures of the colonies (Fig. 6). For the NQS toggle, we observed that the colonies indeed showed vertical segregation, with cells at the top of the colony dome in the blue, and cells at the bottom in the yellow state (Fig. 6C, F). We also sliced the colonies vertically and imaged their cross-sections using confocal microscopy (see Methods). The slices confirmed that cells in the blue state occupied the top of NQS colonies, while cells in the yellow state occupied the bottom, as predicted by the model (Fig. 6F). In contrast, confocal images of QS toggle colonies displayed radial separation between the states, again confirming the predictions of the model, and in agreement with our earlier analysis of experimental findings (Fig. 6A, E). Depending on the aTc concentration, the QS toggle cells at the top of the colony remained yellow, and blue cells only appeared in the outer part of the colony. Imaging of colony slices also confirmed this observation (Fig. 6E).

**Figure 6:**
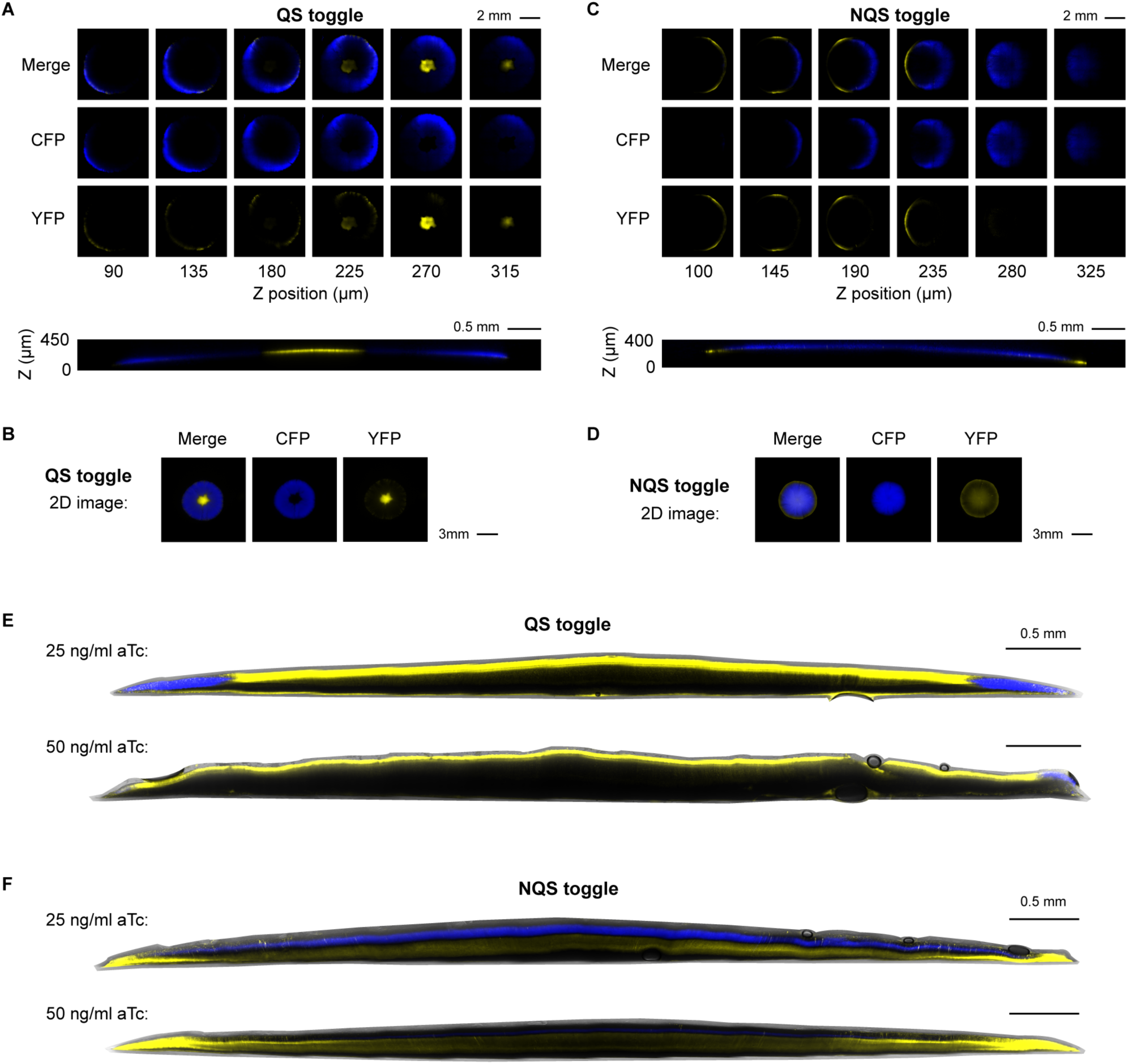
Three-dimensional view of QS and NQS colonies. **A)** QS toggle layers from top to bottom of colony (left to right). Orthogonal view of the QS toggle frames is shown below. Colony is not more than 250 µm tall. **B)** 2D image of the same colony in (A) for comparison. Separate fluorescence channels are shown. **C)** NQS toggle layers from top to bottom of colony (left to right). Orthogonal view of the NQS toggle frames is shown below. **D)** 2D image of the same colony in (C) for comparison. Separate fluorescence channels are shown. **E)** Slices of different QS colonies grown in either 25 or 50 ng/ml aTc. **F)** Slices of different NQS colonies grown in either 25 or 50 ng/ml aTc.

We found that ring formation in the QS toggle exhibited some variability and randomness causing imperfect radial symmetry of the outer blue rings. Despite this variability, the difference in color segregation between QS and NQS colonies was statistically significant (see Fig. 4). Occasionally, we also observed the occurrence of multiple blue rings: *i*.*e*. the formation of internal and external blue rings in QS toggle colonies (Fig. 7 and S5). In addition, patches of blue cells sometimes failed to form a full ring (Fig. S5). These observations suggested that external or internal fluctuations could play a role in determining the observed spatial patterns. Noise can cause jumps between the stable states of a bistable system. However, we expect such switches to be localized in the absence of a mechanism that can synchronize the state across the colony. Thus, we concluded that external fluctuations that affect all or most of the colony were more likely to drive the emergence of complex spatial patterns such as multiple rings. Furthermore, when the QS toggle was grown in a different medium (EZ rich defined medium instead of LB agar), the outer blue ring pattern was disrupted. At times, we observed blue cells emerging from the center of the colony at 100 ng/ml aTc. These patterns resembled those in the NQS toggle in LB agar (Fig. 7 and S6).

**Figure 7:**
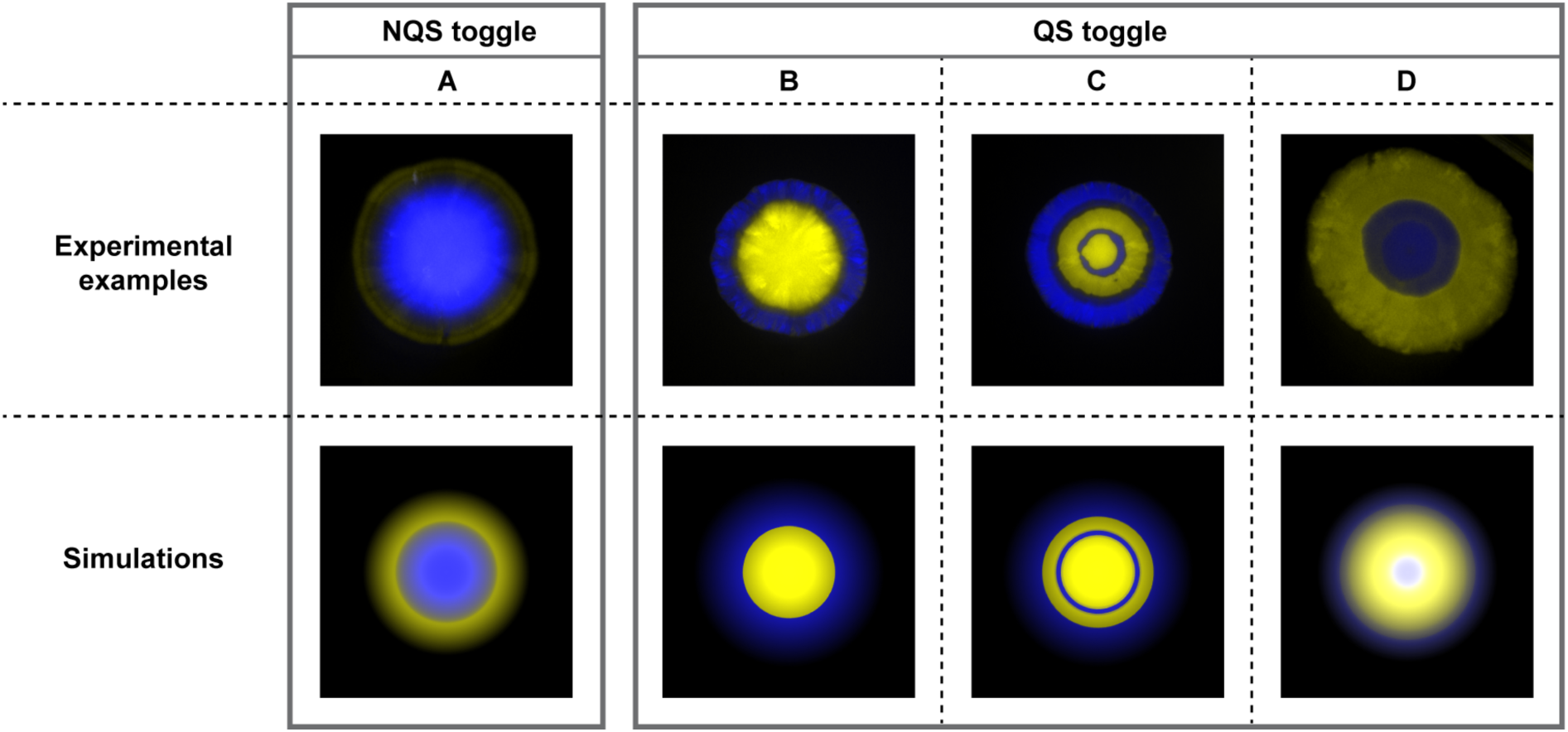
Four types of patterns obtained from experiments (top) and mathematical simulations (bottom). **A)** NQS toggle colony pattern, with vertical segregation of states and blue emergence at the center of the colony. **B)** QS toggle colony pattern, with radial segregation of states, and blue emergence at the edge of the colony, forming an external blue ring. **C)** Alternative QS toggle pattern, with radial segregation of states and multiple blue rings (internal and external). **D)** Inverted QS toggle pattern, where radial segregation of states and blue emergence at the center occur only in specific conditions (see SI).

We were able to obtain similar patterns in numerical simulations by changing parameter values, or by including extrinsic perturbations in the model we described above (Fig. 7 and STAR methods). In particular, we found that extrinsic fluctuations and growth conditions can change effective promoter strength, resulting in the formation of multiple rings and an inverted pattern in the QS system, respectively. This suggests that the complex spatial patterns we observed in some experiments can be explained using the same mechanisms underlying the predominant, single ring patterns. The variability of multi-ring patterns, along with the high dimensionality of the model parameter space makes it difficult to examine such patterns systematically. A full understanding of how these complex patterns emerge, and how they can be controlled will thus require the development of new experimental approaches.

## Discussion

Patterns, and especially ring-like patterns, formed by synthetic gene circuits are not new. Previous studies have utilized various methods to create patterns, such as internal genetic oscillations (Riglar et al., 2019), scale-invariant intracellular signaling (Cao et al., 2016), or mechanical interactions between cell types (Xiong et al., 2020). Here we show that toggle switches can also create patterns in colonies based on how they shape the cells’ responses to a morphogen. Specifically, the classical NQS toggle creates a vertically segregated pattern while the QS toggle leads to a segregated ring structure. We also developed a PDE model that captured both behaviors. An analysis of the model suggests that in the NQS case sequestration of signals and geometry of the colony lead to the formation of a morphogen gradient inside the colony, creating top-down segregation. In contrast, the model suggests that fast cell growth at the periphery of QS toggle colonies and the QS signal gradient inside the colonies lead to the emergence of an outer blue ring. The difference is that, as the aTc level decreases in the QS toggle, signaling between cells in the yellow core helps to lock them in the same state. However, in the NQS toggle, the blue state becomes monostable in the absence of aTc and signals from other cells. As the aTc concentration is lowest at the top of the colony, a wave of transitions from the yellow to the blue state propagates from the colony’s top downward. Our mathematical analysis implies bistability is determined by the external morphogen concentration and the effective promoter strength for each repressor, which is, in turn, determined by protein production, proteolytic degradation, dilution, and QS signals.

Our mathematical model provides further insight into the mechanisms behind the formation of patterns. When modeling pattern formation on the time scale of hours, one must account for expanding colony size. As suggested by Warren et al. (2019), an actively growing layer at the bottom of the colony drives the ‘establishment phase’ (14h – 24h). After 24 hours, the colony enters a ‘flattening phase’, during which vertical growth slows while radial growth remains linear. In our model, the characteristic length scale of signal diffusion is much larger than the size of a colony, and thus details of the colony expansion mechanisms do not have a large impact. We thus assumed that model colonies grow linearly in both vertical and radial directions. The location of newly born cells, on the other hand, does matter. As new cells inherit the state of their parent cells, where the effective promoter strengths of the newly born cells land in the bistable region determines the emergent patterns in the colony.

It should be noted that our model does not precisely predict the full behaviors of the colonies. Our experiments showed that fluorescence intensities were lower in the bulk of many colonies and higher nearer the surface (Fig. 6E). This indicates that the colonies are more complex than our model suggests. Yet, our model still provides qualitative insight into the formation of the patterns and their symmetries. Further experimental studies on *E. coli* colony growth, structure, and metabolism, especially at larger colony sizes, will allow for improvement of the mathematical model and a better understanding of pattern formation in microbial colonies.

Our work indicates that the observed spatial patterns are dependent not just on the underlying genetic circuitry, but also on growth conditions and cellular metabolism. Our mathematical model allows us to identify which patterns are allowed by the system and the mechanisms that generate them. The presence or absence of intercellular communication, for instance, is not the determinant of a single pattern. Rather, the resultant pattern is determined by multiple factors, including morphogen concentration, promoter strengths, and intercellular signaling. Nevertheless, the presence of intercellular signaling allows one to actively control or shape the balance by implementing more spatial features, such as signal degradation, by differentiated cells or external flux. These patterns shed light into how synthetic multicellularity can be created in bacteria and provide a further step toward the creation of large scale programmable synthetic multicellular systems.

## Acknowledgements

B.F.M. acknowledges support from Brazilian Coordination for the Improvement of Higher Education Personnel (CAPES), through the Science without Borders (SwB) fellowship. G.F. and K.J. acknowledges support from NSF (1936770). K.J. acknowledges support from MCB-1936770. M.R.B. acknowledges support from the Welch Foundation (C-1729), the National Institutes of Health (R01GM144959), and the NSF (MCB-1936774). E.D.S. acknowledges support from grants NSF/DMS-2052455 and AFOSR FA9550-21-1-0289. The authors also acknowledge the use of resources of the Shared Equipment Authority at Rice University for this work. The computational work was completed in part with resources provided by the Research Computing Data Core at the University of Houston.

## Author contributions

Conceptualization, B.F.M., G.F., E.D.S., K.J., M.R.B.; methodology, B.F.M., G.F., E.D.S., K.J., M.R.B; validation, B.F.M., M.R.B.; data curation, B.F.M.; investigation, B.F.M., G.F.; formal analysis, B.F.M., G.F; visualization, B.F.M., G.F.; writing – original draft, B.F.M., G.F., K.J., M.R.B.; writing – review & editing, B.F.M., G.F., E.D.S., K.J., M.R.B.; software, G.F.; project administration, B.F.M., G.F., K.J., M.R.B.; funding acquisition, B.F.M., K.J., M.R.B.; supervision, K.J., M.R.B.

## Main figure titles and legends

(In text)

## STAR Methods

### Methods

#### Plasmids and strains

We constructed plasmids with either PCR-based, restriction enzyme cloning, or Golden gate assembly methods. QS and NQS toggles, as well as all other tested versions, are composed with 3 plasmids each, providing resistance to kanamycin, chloramphenicol and spectinomycin. A list of all plasmids employed is provided in Table S1. For this study, we used the CY027 *E. coli* strain, a BW25113 derivative (Chen et al., 2015). This strain has the *lacI, araC* and *sdiA* genes knocked out, and constitutive *cinR* and *rhlR* knocked-in to its genome to enable QS communication (*ΔlacI ΔaraC ΔsdiA Ptrc*-cinR Ptrc*-rhlR*).

#### Plate reader experiments

From single colonies, we inoculated cells containing the appropriate plasmids into 5 mL LB with antibiotics (50 *μ*g/mL kanamycin, 34 *μ*g/mL chloramphenicol, 50 *μ*g/mL spectinomycin) for overnight growth at 37°C in a shaker (250 rpm). Then, we diluted the culture 1:100 in minimal M9CA broth (Teknova) with antibiotics, and we grew these cells for 2 hours in a 37°C shaker (250 rpm). Meanwhile, we prepared 96-well round bottom plates with minimal M9CA broth, antibiotics, and applicable inducers (IPTG and aTc) with a 2-fold final concentration of a volume of 100 *μ*L per well. After the 2-hour growth, we added 100 *μ*L of cell outgrowth to each well (1:1). We incubated the plates at 37°C, shaking at 800 rpm. After 2 hours, we read each plate in a Tecan Infinite M1000 for growth (OD, 600 nm), YFP fluorescence (ex, 514 nm; em, 527 nm), and CFP fluorescence (ex, 433 nm; em, 475 nm). We used cells without plasmids to measure background auto-fluorescence. The results shown are reported as (fluorescence-background)/OD_600_.

#### Flow cytometry

We inoculated cells from single colonies into 5 mL LB with antibiotics and either 0.5 mM IPTG (pre-IPTG) or 50 ng/ml aTc (pre-aTc) for the overnight growth at 37°C in a shaker (250 rpm). Next, we prepared 96-well round bottom plates with minimal M9CA broth, antibiotics, and applicable inducers (IPTG and aTc) with a 2-fold final concentration at a volume of 100 *μ*L per well. We diluted the overnight cultures 1:50 in minimal M9CA broth with antibiotics, and we added 100 *μ*L of this cell dilution to each well (1:1, final cell dilution of 1:100). We incubated the plates at 37°C, shaking at 650 rpm. After 3 hours (and 9 or 12 hours for stationary phase tests), we kept the plates on ice for at least 10 min. Then, we added 25 *μ*L of each well to a tube with 475 *μ*L of 1x PBS (5% dilution), and vortexed the tube. We analyzed each tube with the Sony SH800S Cell Sorter. We used filters for mCFP and EYFP. Due to overlap in their fluorescence spectra, we used single fluorescence controls and the Sony software calculated compensations for each fluorophore. All CFP and YFP values shown here are compensated. For each sample, we recorded 10,000 events. We exported and analyzed the acquired data with FlowJo. We manually created the fluorescence gates. The common existence of OFF cells among the circuits (i.e., cells that are expressing neither CFP nor YFP) generated the need for an OFF gate. We created blank controls for each circuit by transforming the circuit plasmids with an empty reporter plasmid instead of the regular CFP/YFP one. We used these circuits to determine the OFF gate. Next, we created CFP+ and YFP+ gates by drawing a diagonal line from top right to bottom left, until it reaches the OFF gate. CFP+ gate is at the upper left of this line, while YFP+ is at the bottom right. We exported geometric mean values and population composition based on these gates from FlowJo and plotted with Matlab. For the confocal tests, we also used the Sony SH800S to sort single cells into small petri dishes (60 mm) containing LB agar. We attached the dishes to the 96-well-plate stage for sorting, and later put them into 37°C incubator for growth into single colonies.

#### Colony tests

We prepared LB agar plates with antibiotics (50 *μ*g/mL kanamycin, 34 *μ*g/mL chloramphenicol, 50 *μ*g/mL spectinomycin), and aTc (12.5, 25, 50 or 100 ng/ml). We inoculated cells from single colonies into 5 mL LB with antibiotics and 0.0625 mM IPTG for overnight growth at 37°C in a shaker (250 rpm). Next, we diluted the culture 1:100 in 4 mL LB with antibiotics and 0.0625 mM IPTG, and we grew the culture at 37°C shaker (250 rpm) until it reached an OD_600_ between 0.7-0.8 (approximately 2 hours). We then diluted the culture with LB to reach an OD_600_ range between 0.35-0.4. Then, we used 1 mL of the diluted outgrowth for serial dilutions until we achieved 1:10,000 and 1:100,000 (outgrowth:LB) ratios. We put plates to warm up in 37°C incubator for at least an hour before plating. Then, we plated both 1:10,000 and 1:100,000 dilutions into 2 equal sets of plates, with 12 glass beads per plate. We wrapped plates in foil to avoid light exposure of aTc, and we put them in a 37°C incubator. We took the plates at 24, 28, 32, 44, 52, 76 and 100h post-plating to be imaged in a low-magnification microscope (see 2D imaging).

#### HPLC test

To quantify aTc concentration from LB agar fragments, we modified the method from (Halling-Sørensen et al., 2002). We used an Agilent 1220 Infinity LC instrument, and a C-18 chromatographic column (Aeris 3.6 *μ*m Peptide XB-C18 100A LC column 250 × 4.6 mm). To start, we directly tested aTc in the following amounts: 5 *μ*g, 2.5 *μ*g, 1.25 *μ*g, 0.625 *μ*g, 0.3125 *μ*g, 0,15625 *μ*g. From the area under peak, we generated a calibration curve and an equation for aTc quantification for the following tests (Fig. S3A). We prepared triplicates of 13 mL LB agar with aTc at final concentration of 100 *μ*g/mL. Then, we poured 6 mL of each triplicate into two 24-well plates (1 mL/well). We prepared a total of 36 wells (triplicates for 6 reads in 2 plates). To measure the aTc, we transferred the solidified media from each well to separate tubes, and we diluted each with 9 mL of water. The tubes were kept in the dark and at RT for 2 hours. Then, we centrifugated the tubes and filter sterilized the liquid. First, we measured triplicate samples (100 *μ*L injection each) in HPLC before any actual incubation (day 0). Then, we incubated one plate for 48 hours at 37°C (the one without a set of triplicates from day 0 read), while we incubated the other plate at 4°C to recreate the two possible pre-treatments, as described in Colony tests section. After the first 48 hours, we also incubated the 4°C plate at 37°C, except for a set of triplicates that was kept at 4°C until the end of the experiment. We continued the incubation for a total of 6 days to recapitulate the actual experimental setup (see Colony tests). Each day starting at day 2, we removed a set of triplicates from each plate, then, treated and injected in the HPLC as explained above. Finally, we quantified aTc by measuring the area under the peak for each run.

#### 2D imaging

We imaged plates with colonies with a stereo microscope (Nikon SMZ800), YFP and CFP fluorescence filters (Chroma #39003 and #39001, respectively), and NIS-Elements software (Nikon). QS and NQS toggles are very different circuits, and therefore, have different fluorescence intensities. For this reason, we used distinct settings for each circuit. We imaged QS toggle colonies with an exposure time of 300 ms for YFP and CFP. While NQS toggle had lower intensities, we used an exposure time of 1 s for both YFP and CFP. We manually imaged each frame for CFP, YFP and bright field to form a multichannel image. Therefore, not all colonies were captured from all plates. Whenever there were many colonies, we generally picked 4 frames per plate to cover as many well-separated colonies as possible. We either did not image or did not analyze colonies that were in contact with other colonies. We exported each multichannel image into 3 separate tiff files. We assembled colony images with Fiji (ImageJ). We used identical minimum and maximum fluorescence values among images in the same figure to enable direct comparison, but different values across circuits. For confocal tests, we also imaged colonies with the stereo microscope before any agarose was added. In such images, we used the same exposure time settings for YFP and CFP for QS and NQS toggles (1 s for YFP and 300 ms for CFP). In Fiji, we did not use the same adjusted minimum and maximum fluorescence values for QS and NQS toggles, making these images not directly comparable in respect to their fluorescence intensities.

#### Confocal imaging

We sorted single cells into LB agar plates (see Flow cytometry), containing antibiotics and aTc (25 ng/ml or 50 ng/ml). We incubated the plates at 37°C, wrapped in foil, for different times, ranging from 32 to 72 hours. After growth, we covered each colony with approximately 1 mL of an autoclaved 1% agarose solution. For the full colony images, we cut out pieces of agar with agarose-covered colonies, we placed them upside down (with agarose on bottom) on glass coverslips and took them to the Nikon A1 Confocal microscope. For the perpendicular cross-section, we manually cut the pieces in the middle with a blade. We placed the middle part into a glass coverslip for imaging on a Nikon A1 Confocal microscope. In the confocal, we used 405 nm and 488 nm lasers for excitation of CFP and YFP, respectively. We acquired images with 10x (for full colony Z-stacks) or 20x objective (for slices), resonant scanner, 1 A.U., with denoise capture mode. For Z-stacks, steps varied from 5.1 to 15 *μ*m distance, for a total of 200-500 *μ*m coverage within the Z axis. We exported the data into separate tiff files. We made videos with Fiji (Image J). We used identical minimum and maximum fluorescence values among images from the same circuit replicates to enable direct comparison, except for figures of slices. We also captured colonies with stereo microscope (2D imaging) prior to agarose solution addition for comparison.

### Data analysis

We performed all data analyses in Matlab. First, we manually selected all colonies each frame, and we took the average YFP and CFP fluorescence of four radii (one per direction: N, S, E, W) per colony. We used the bright field for mask creation and detection of colonies. We selected regions without colonies for background, which we subtracted from initial intensities. We saved all data from all colonies in the same plate in a matrix per time point. We determined the yellow width by measuring the distance from the center of the colony to the threshold point, in which yellow fluorescence first reaches below 30% of its maximum. We then normalized this distance by dividing with the total colony length (radius), giving results between 0 and 1. To obtain this measure we made several assumptions: if the maximum YFP value was below a certain arbitrary threshold of 200, the yellow color was considered absent, and the normalized yellow width was 1 divided by total length (close to zero). If the maximum CFP value was below an arbitrary threshold of 300, the blue color was considered absent, and the normalized yellow width was 1. When the maximum YFP intensity at a particular pixel also had CFP above a very high second threshold of 2000, this probably represented a random patch of yellow cells between blue cells, and we assumed yellow was absent for purposes of width calculation. Lastly, when blue was absent, and YFP showed small variation between maximum and minimum (below 170), the colony was considered completely yellow, we selected the full colony as the width, instead of the highest value. We picked all these arbitrary threshold values for CFP and YFP through observation. We used them equally for all colonies of both circuits. Furthermore, we normalized each colony’s original fluorescence data by its maximum to allow for color overlap analysis. We plotted each pixel from the center to the edge of the colony by its normalized YFP and CFP fluorescence, and they were only classified as overlapping when both YFP and CFP values were above the threshold of 0.3. Some colonies were found to have only one color throughout the entire radius, for example, some of all-yellow colonies at high aTc concentration. Therefore, such colonies do not show any overlap, and can skew the data. To remove single color data (Fig. S4), we selected only colonies that had both colors present for at least 25% of pixels. Finally, we compared all data from QS colonies to NQS. Both yellow width and overlap data showed a non-normal distribution, thus, we performed non-parametric statistical tests. We analyzed comparisons among 4 different aTc concentrations of the same circuit with Kruskal-Wallis test (non-parametric equivalent to one-way ANOVA), followed by multiple comparison test (Tukey-Kramer). While we analyzed direct comparison among 2 sets of data with Mann-Whitney test (non-parametric equivalent to Student’s t-test).

## Mathematical model

### Section I – Bistability in Well-mixed Environment/Liquid Culture

In the following we describe the set of equations modeling the concentrations of LacI and TetR within the cells in the population. We assume that these concentrations are approximately proportional to the fluorescent signals which are measured experimentally and use this assumption to compare experimental and modeling results.

#### 1. Single-cell Model Derivation

We let x(t) and y(t) represent the concentration of LacI and TetR, respectively. Assuming that the intracellular concentrations of LacI and TetR are proportional the concentrations of the fluorescent proteins YFP and CFP, respectively, the quantities x(t) and y(t) are then directly related to the experimentally measured fluorescence. We therefore compare the evolution x(t) and y(t) predicted by our model to the experimentally measured fluorescence signals. The production of LacI is regulated by a promoter, which can be activated by QS signaling molecules (C14-HSL bound to CinR), and repressed by TetR. Similarly, production of TetR is activated by the QS signal (C4-HSL bound to RhlR), and repressed by LacI. We denote by *g*(*t*) and *h*(*t*) the concentration of QS signals C14-HSL and C4-HSL at time *t*, respectively. We also assume that LacI and TetR can be produced at maximum rates *a*_1_ and *a*_2_, and are degraded with rate constants *γ*_1_ and *γ*_2_, in addition to constant dilution at rate *γ*_*d*_ due to cell growth. We can then model the dynamics of the QS toggle using the following system of ODEs describing the intracellular concentrations of LacI and TetR in one of the cells in the colony (Gardner et al., 2000b; Nordholt et al., 2017; Zong et al., 2018):

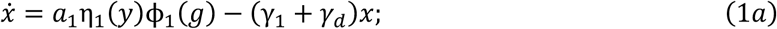

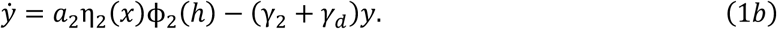

The repression of the promoter is represented by Hill functions *η*_1_ and *η*_2_, while activation is represented by Hill functions ϕ_1_ and ϕ_2_. In particular, we set

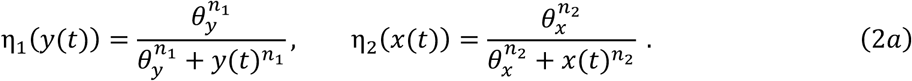

and

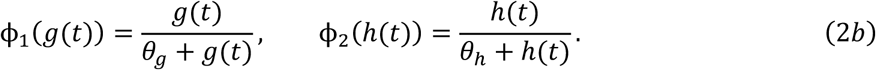

Here, the choice of Hill coefficients is based on the number of monomers in the activator (1 for C14 and C4) and repressor (*n*_1_ = 2 for TetR and *n*_2_ = 4 for LacI).

We model the dynamics of the intercellular QS signals, C14-HSL and C4-HSL, whose activation is repressed by TetR and LacI, by the following equations,

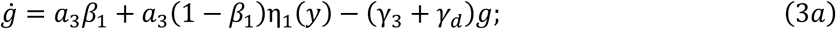

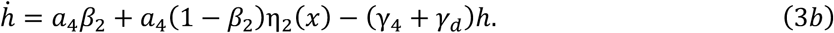

Here, C14-HSL and C4-HSL are produced with maximum rate *a*_3_ and *a*_4_, and degraded at rate *γ*_3_ and *γ*_4_, respectively. Further, let *a*_3_*β*_1_ and *a*_4_*β*_2_ represent the base production rate of the QS signals. Setting ϕ_1_(*g*(*t*)) = 1 and ϕ_2_(*h*(*t*)) = 1 results in equations that describe the NQS toggle.

When exogenous inducers, IPTG, and aTc, which we denote by U and V, are added to the well-mixed liquid culture, the state of the system can be changed. Following experiments, we will assume that IPTG and aTc are provided at some background concentration level, *u*_0_ and *ν*_0_, respectively. De-repression of the production of LacI/TetR is achieved through inducers binding to their corresponding repressors. Let X and Y represent the unbound/free repressor, LacI and TetR, correspondingly. If 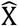 and 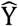 denote the inducer-bound complex the binding activity can be described by the following reaction scheme,

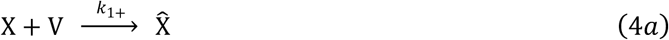

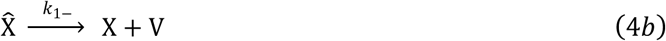

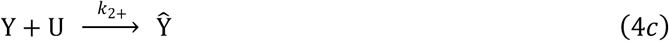

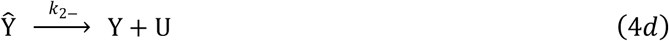

Let *x*_*b*_ (*t*) and *y*_*b*_ (*t*) represent the concentration of bound LacI 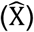 and bound TetR 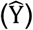, we then have that

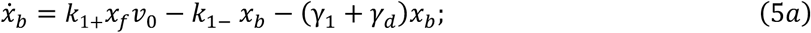

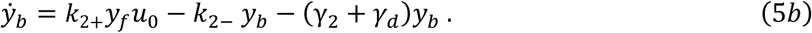

Here, *x*_*f*_ (*t*) and *y*_*f*_ (*t*) represents the free LacI and TetR. By conservation law, we have that *x*(*t*) = *x*_*b*_(*t*) + *x*_*f*_(*t*) and *y*(*t*) = *y*_*b*_(*t*) + *y*_*f*_(*t*).

Hence, the set of equations describing the dynamics of the intracellular concentrations of LacI, TetR, the signaling molecules, as well as the concentrations of bound LacI and TetR is given by,

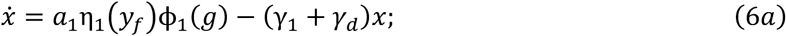

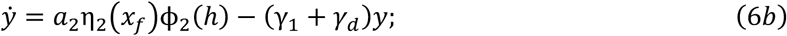

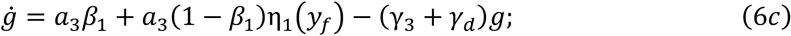

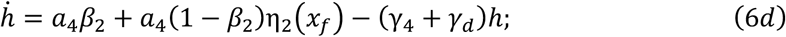

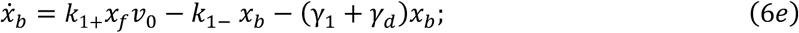

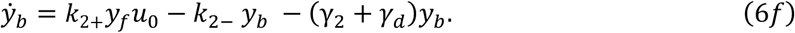

#### 2. Nondimensionalization and Bifurcation Analysis

To simplify analysis, we next assume that *γ*_1_ = *γ*_2_ = *γ*. Defining the rescaled time *τ* = (*γ* + *γ*_*d*_) *t*, we can nondimensionalize each variable using its corresponding threshold to obtain the following nondimensionalized QS toggle system,

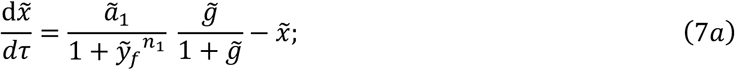

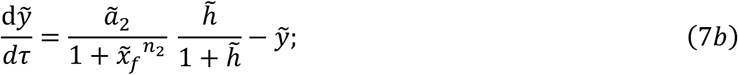

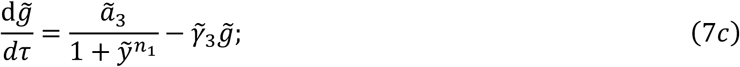

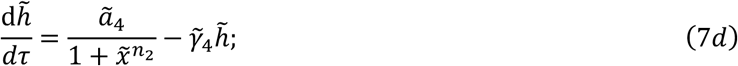

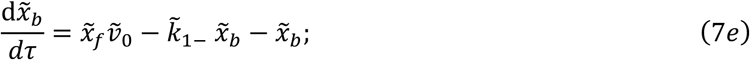

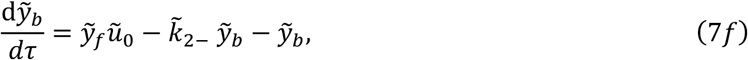

with 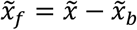 and 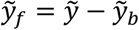. Here the nondimensionalized variables and parameters are,

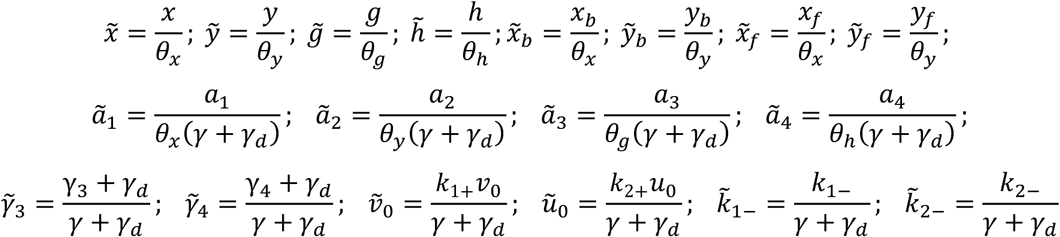

Eq. (7) describes the NQS toggle system after removing (7c-d) and setting 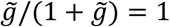 and 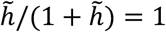 in (7a-b). The nondimensionalization shows that, in the NQS toggle system, the strength of the repressor, *ã*_1_ and *ã*_2_, is determined by a balance between production, degradation, dilution and the repression threshold. In the QS system, the effective repressor strength is modified by the QS signal profile. That is,

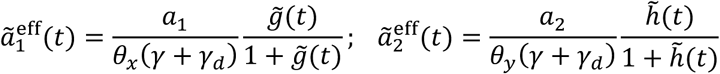

To understand how the different parameters impact the equilibria of the system, we first obtain the equilibria by setting the derivatives in Eq. (7) to 0. Denoting the equilibrium values of the different dynamical variables by a star, we have that, 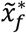 and 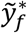 satisfy,

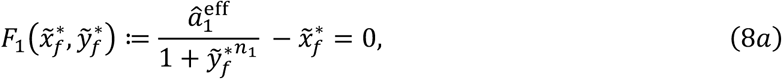

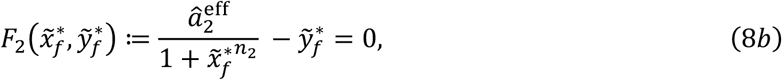

With

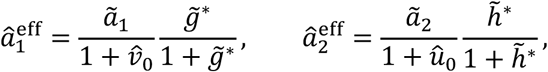

and

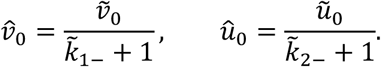

Let *ℒ* ⊂ ℝ^2^ represents the bifurcation curve in the 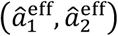 parameter space, that is the curve in parameter space at which the behavior of the system changes qualitatively. Thus, crossing the bifurcation curve leads to a change in the number of equilibria and/or their stability (Kuznetsov, 2004). Except for the steady-state condition described in equations (8a-b), points on the bifurcation curve *ℒ* additionally must satisfy a matching slope condition (Kuznetsov, 2004). That is,

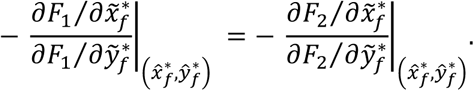

After some algebra, the above condition can be simplified as following,

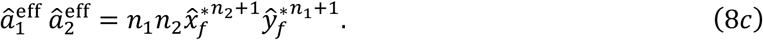

Solving system (8) numerically gives us the cusp bifurcation curve *ℒ* in the 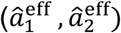 space, as shown in Figure 2E. Nullclines plotted in Figure 2F, G are given by solving system (8a-b) for 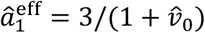 and 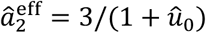.

### Section II – Spatial-temporal Dynamics of the NQS and QS colony

Using the model of the intracellular dynamics developed above, we next describe a model of the spatiotemporal dynamics of the corresponding quantities in a growing bacterial colony.

#### 1. Domain

The domain, Ω ⊆ ℝ^3^, on which we define the model consists of two different subdomains, Ω = Ω_1_ ∪ Ω_2_, with Ω_1_ representing the part of the domain occupied by the bacterial colony, and Ω_2_ the part occupied by the agar. The agar plate takes the shape of a cylinder, and we assume that the colony takes shape of a cone (Warren et al., 2019b), as show in Fig. S7A. Assuming radial symmetry, we can reduce the 3D domain to a 2D slice.

There are two different types of boundaries: The inner boundary, represented by *∂Ω*^in^, and the outer boundary, represented by *∂Ω*. The inner boundary is the interface between the colony subdomain, *Ω*_1_, and the agar subdomain, *Ω*_2_. The outer boundary is the union of the colony-air interface and the agar-plate interface.

The colony is observed experimentally over tens of hours during which it can grow substantially. We therefore include colony growth in the model. Previously, Warren et al. (2019) have identified three phases of colony expansion: The initial monolayer phase (0-13 h); the establishment phase (14-24 h); the flattening phase (24+ h). In the establishment phase, growth in both height and radius is linear, as cells predominantly divide in the active growing region consisting of a thin disk at the bottom of the colony of approximate height H_AGL_ = 10 *μm*. During the flattening phase, radial growth is still linear while the increase in height slows.

In our case, the observed pattern in both the NQS and QS colony is driven by diffusive signals. Since the size the colony is much smaller than the characteristic length scale of signal diffusion during the experiments, details quantitative description on how the colony grows vertically do not matter much. Therefore, for simplicity we assume that the colony expands linearly in both radius and height, with a thin actively growing layer on the bottom. These assumptions together imply a fixed aspect ratio in colony height vs radius. In our simulation, the continuous growth of the colony is discretized by adding slabs of uniform heights and linearly increasing radius every unit time. In particular, in experiment the radius of the colony by the end of 100 hours is approximately 4 *mm*, giving a linear radius growth rate of *ν*_*r*_ = 40 *μm*/*h*. Moreover, the cross-sectional images of the colony show that at different times of colony expansion, the height to radius ratio ranges from 1: 7 to 1: 12. For simplicity, we assume a fixed ratio of the height to base in the triangular slice to be 1: 10, which leads to a linear height growth rate of *ν*_2_ = 4 *μm*/*h*.

**Supplement Figure 7:**
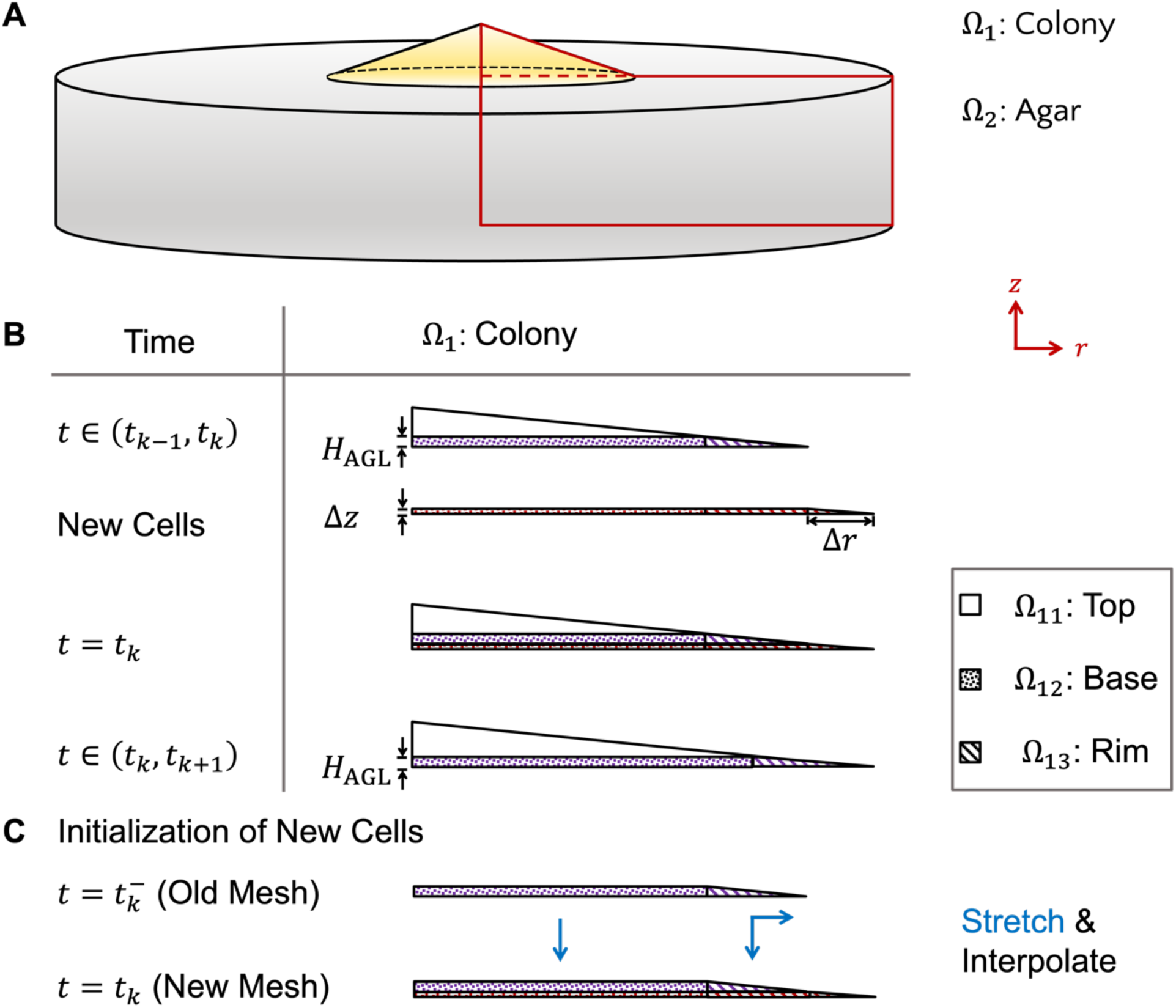
Schematics of the 3D domain and the discretization of the colony expansion used in numerical simulations. **A)** We assume a cone shaped growing colony sits on top of the cylindrical agar pad. The red curves outline a 2D slice from the 3D domain, in the radial (*r*) and height (*z*) direction. Assuming radial symmetry, which is consistent with experimental observations, we used this 2D slice as the domain for the model. The solid line represents the outer boundary (*∂Ω*) while the dashed line represents the inner boundary (*∂Ω*^in^). **B)** Growth is modeled by updating the colony’s shape at even increments in time, *Δt*. At the end of each subinterval, the domain of the colony, *Ω*_1_, is increased by adding a rectangle of height *Δz* and of width equal to that of the colony, and an adjoining right triangle of height *Δz* with base Δ*r*. Here, we discretize time into intervals (*t*_−1_, *t*_+_), with *t*_*k*_ = *k* · Δ*t*. **C)** After every increment of time, a new mesh is generated for the expanded colony. The new nodes at the Top and Rim region are initialized by interpolating the solution of the corresponding region from the stretched old mesh.

As shown in Fig. S7B, based on growth assumption at different locations, we divide the colony subdomain *Ω*_1_ into three different regions, top (*Ω*_11_), base (*Ω*_12_), and rim (*Ω*_13_). In particular, the triangular top, defined as *Ω*_11_ = {(*r, z*) ∈ Ω_1_|*z* ≥ *H*_AGL_}, doesn’t grow. Let *R*_*k*_ represents the radial length of the colony at the end of time intervals (*t*_*k*71_, *t*_*k*_). The rectangular base, defined as *Ω*_12_ = {(*r, z*) ∈ Ω_1_|*r* ≤ *R*_*k*_, *z* ≤ *H*_AGL_}, grows linearly only in the vertical direction. The triangular rim, defined as *Ω*_13_ = {(*r, z*) ∈ Ω_1_|*r* ≥ *R*_*k*_}, grows linearly in both the vertical and radial direction.

Let *T* be the time it takes for the colony to grow *H*_AGL_ in the vertical direction. Here, *T* = *H*_AGL_/*ν*_2_ = 150 *min*. Let 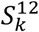 represents the area of the top region, *Ω*_12_, at the end of time interval (*t*_*k*71_, *t*_*k*_), where 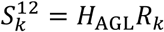. Then we have that

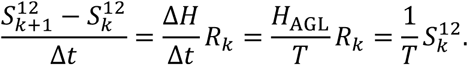

Similarly, let 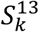 represents the area of the rim region, *Ω*_13_, at the end of time interval (*t*_*k*71_, *t*_*k*_), where 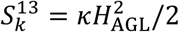, with *κ* represent the fixed radius-height ratio of the colony. Then we have

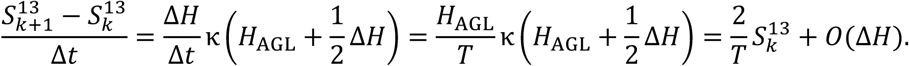

Therefore, we can approximate the expansion factor of the rim region, *Ω*_13_, by 2/*T*, which is twice as fast as the expansion rate base region, *Ω*_12_. Let *γ*_*d*_ = 1/*T*, we then get that the following chemical dilution rate from colony expansion in the different regions,

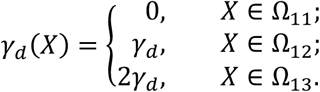

#### 2. PDE Model

Let *u*(*X, t*) represent the aTc concentration at time *t* and location *X* ∈ Ω. The inducer, aTc, is initially supplied in the agar, which then diffuses into the colony and react with TetR in different cells. The reaction occurring within the cells in the colony can be modeled using Eqs. (4*c*, *d*). We assume that when the aTc-TetR complex, *Ŷ*, is being degraded by the enzyme ClpXP, and that a portion *α* ∈ [0,1] of the aTc returns to the cell from the complex (Nevozhay et al., 2009). That is,

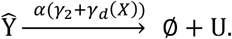

Putting everything together, for the concentrations of,

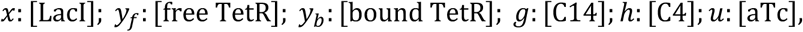

with *y* = *y*_*f*_ + *y*_*b*_, and the superscript represent different subdomain, we obtain a diffusion-reaction model we describe next.

In the colony domain, X ∈ Ω_1_, the various concentrations evolve according to,

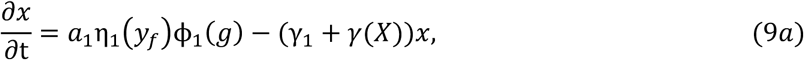

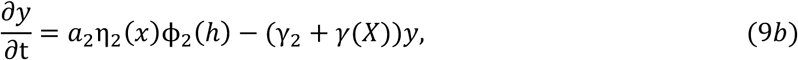

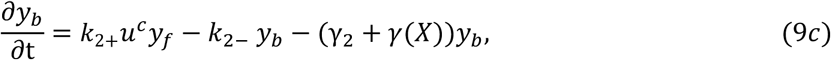

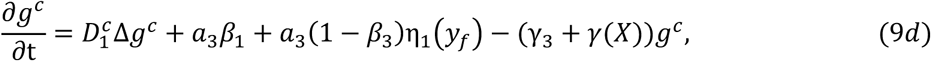

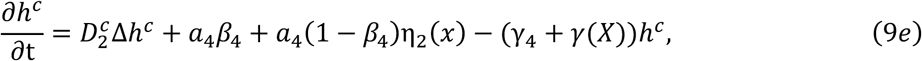

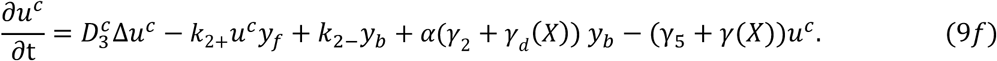

In the agar domain, Ω_2_, we have

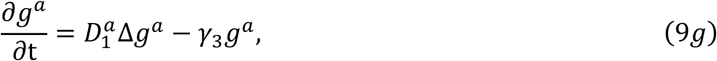

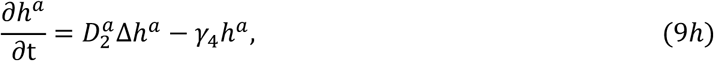

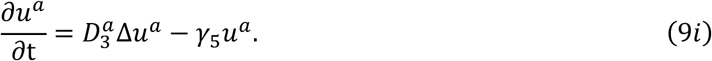

Here 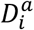 denotes the signal diffusion coefficient in agar, for different chemicals C14, C4 and aTc indexed by *i* = 1,2,3 respectively. Due to the crowdedness of the colony, we assume that signal diffusion coefficients in the colony, 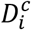, satisfies 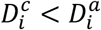. In particular, it has been shown that GFP diffuse 10 times faster in water 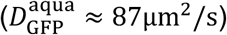 (Swaminathan et al., 1997), compared with in the cytoplasm 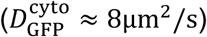 (Elf et al., 2007). Therefore, we set 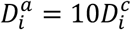. Since aTc has a lower molecular weight than GFP, we assume that the aTc diffusion coefficient in agar is 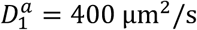. It has been reported that the effective diffusion coefficients of C14 and C4 are *D*_C14_ ≈ 83µm^2^/s and *D*_C4_ ≈ 1810µm^2^/s (Karig et al., 2018). Here we set the diffusion coefficients in agar to 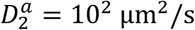 and 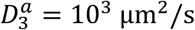, for C14 and C4, respectively.

For the diffusible chemicals, *C* = *g, h*, or *u*, we also we the following inner and outer boundary conditions.

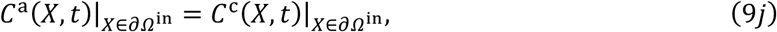

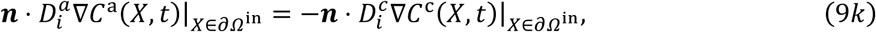

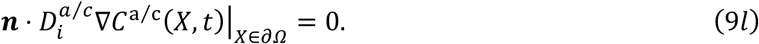

We used the radial symmetry of the 3D domain to reduce it to an equivalent 2D model.

Let *r* and *z* represent the independent variables denotes the radius and height coordinate inside the colony. We then have that the Laplacian operator acts on *C* = *g, h*, or *u* as

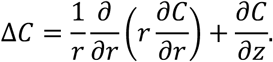

### Parameters and Simulation

We assume the tagged degradation gives a half-life of 7 min for intercellular species (Chen et al., 2015). This leads to degradation rates of LacI and TetR of *γ* = *γ*_1_ = *γ*_2_ = ln(2) /7 min^−1^. It has been reported that the degradation rate of AHL ranges from zero (no detectable degradation at 32 hours) to a half-life of 8 hours depending on environmental factors such as temperature and pH value (Politi et al., 2014). For our experimental condition, we assume that the half-life of C14 and C4 is 24 hours, corresponding to degradation rates of *γ*_3_ = *γ*_4_ = ln(2) /24 *h*^−1^. The production rates of each species, *a*_*i*_, with *i* = 1,. ., 4 along with their corresponding threshold of activation (EC50)/repression (IC50) parameters *θ*_*j*_, with *j* = {*x, y, g, h*}, were chosen so that the nondimensionalized parameters, *ã*_*i*_, give a match with experimentally observed patterns. We also set the basal production level of the AHL signals to *β*_1_ = 0.2 and *β*_1_ = 0.5, for C14 and C4 respectively. It has been reported that aTc bind with TetR at rate, *k*_+_ = 0.06/nM/min (Nevozhay et al., 2009). As shown in Fig 1E and G, aTc can induce the yellow state at a concentration of 1 ng/ml in the NQS, and at approximately 10 ng/ml in the QS case. Therefore, we set *k*_<_ = *k* · 0.06 nM^−1^min^−1^, with *k* = 1 in the NQS case and *k* = 10 in the QS case. We set the unbinding rate to *k*_−_ = *k*_+_/*k*_*A*_, with association constant *k*_*A*_ = 10 nM^−1^ (Kintrup et al., 2000). When the aTc-TetR complex is degraded by ClpXP, we assume that *α* = 0.8 portion of the aTc in the complex returns back to the cell. All parameters used in Figure 7C,D are the same as those used in Figure 7B except the following: in Figure 7C, a pulse of *γ* = ln(2) /5.178 min^−1^ between *t* = 23.25 hr and *t* = 25.575 hr is applied; in Figure 7D, repressor strengths are reduced to *a*_1_ = 80 nM/min and *a*_2_ = 120 nM/min.

We simulate the PDE-ODE model, described by Eqs. (9) using MATLAB. To reduce interpolation error, we took the following approach: First, over *δt* = 1min time intervals, the PDEs were solved using the Partial Differential Equation Toolbox and the ODEs were solved using ode15s at each mesh point, using the solution from the previous time step as initial conditions. Second, the mesh for subdomain Ω_11_, the top of the colony, was kept the same between each update of the colony size at every Δ*t* = 15 min. Meanwhile, at each colony growth update time point *t* = *t*_*k*_, as shown in Figure S1B, mesh points in the bottom layer (Ω_12_ and Ω_13_) were initialized by interpolating the solution at 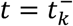 with the coordinates of the mesh points linearly stretched, as shown in Figure S1C.

## Supplemental information titles and legends

**Supplemental Figure 1: Individual behavior of QS and NQS toggle cells when treated with a single inducer for different duration of times.**

Flow cytometry data of QS and NQS toggle cells that were pre-induced with either IPTG or aTc. Each dot is a single cell classified within a gate. Gates were determined with single color and double negative controls. Dashed lines in each plot represent the boundaries between the three distinct gates, which represent cellular states: CFP+ (top gate), YFP+ (bottom-right gate), and OFF (bottom-left gate). Background colors in each plot represent which state the majority of cells are in (>50%): blue color indicates mostly CFP+ cells, yellow plots are mostly YFP+, gray plots are mostly OFF, and white plots indicate cells that are present in multiple states (<50% each). **A)** QS toggle cells pre-treated with IPTG and aTc after growth for 3 (top), 9 (middle), and 12 hours (bottom). **B)** NQS toggle cells pre-treated with IPTG and aTc after growth for 3 (top) and 9 hours (bottom) (see Fig. 2).

**Supplemental Figure 2: Induction curves of circuit variations of the QS toggle.**

We obtained the Weaker Blue State by using lower strength promoters for *tetR* and *cfp* genes. We obtained the Weaker QS toggle by using lower strength promoters for the circuit genes, except both QS genes which were kept the same. We obtained the Inverted QS toggle by using the same lower strength promoters but reversing the QS network connected to each state: now, the rhlR/I network activates the yellow state, and cinR/I the blue state. **A, B) I**nduction curves of Weaker Blue State QS toggle with IPTG (A) and aTc (B) in liquid culture. **C, D)** Induction curves of Weaker QS toggle with IPTG (C) and aTc (D) in liquid culture. **E, F)** Induction curves of Weaker QS toggle with IPTG (E) and aTc (F) in liquid culture, in which cells were pre-treated with IPTG. **G, H)** Induction curves of weaker QS toggle with IPTG (G) and aTc (H) in liquid culture, in which cells were pre-treated with aTc. **I, J)** Induction curves of Inverted QS toggle with IPTG (I) and aTc (J) in liquid culture. Lines represent the average fluorescence and error bars represent the standard deviation of 3 technical replicates for at least 3 independent experiments (see Fig. 1).

**Supplemental Figure 3: Incubation at 37°C causes aTc degradation.**

**A)** Yellow ring width measured from QS toggle colonies grown in aTc plates pre-incubated either at 4°C or 37°C for 48 hours prior to plating. Then, we plated and grew cells as shown in Fig. 3A. These values represent yellow ring widths from 76h post-plating. The 37°C plates showed a significant decrease in width in comparison to its 4°C counterparts, except at 12.5 ng/ml (\**p* < 0.01, Mann-Whitney non-parametric test). Data is from 5 independent experiments. Pre-4°C colonies are included in Fig. 3B. **B)** ATc quantification from LB agar extracts (in the absence of cells) with an HPLC. At time 0, aTc was quantified before any incubation. Samples were divided in two groups: pre-incubation for 48 hours at 37°C (bright blue), or 4°C (light blue) to recreate the experimental timeline in (A). On day 2, we incubated both groups at 37°C until day 6 to also recapitulate the experimental setup. As a control (gray), LB agar + aTc samples were kept at 4°C throughout the entire test and measured on day 7. Data represents mean ± standard deviation of 3 independent experiments (see Fig. 3).

**Supplemental Figure 4: Quantification of pixel overlap for QS toggle (light gray) and NQS toggle (dark gray) colonies without single color data.**

We selected only colonies that have both colors present for at least 25% of the radius. We normalized the number of overlapping pixels by the colony radius (total pixels) (*p* < 0.01, Mann-Whitney non-parametric test). Data for each QS toggle test contains over 138 colonies from at least 11 independent experiments, while data for each NQS test contains at least 5 colonies from 2 independent experiments (see Fig. 4).

**Supplemental Figure 5: Multiple blue rings are also observed in LB agar QS colonies.**

**A)** Example of colony obtained from a 50 ng/ml aTc plate, over time. **B, C)** Fluorescence intensity cross-section at 52h and 76h, respectively, shown in (A). Curves are the average fluorescence between 4 radii of the same colony. **D, E)** We plotted each colony pixel from the images at 52h and 76h, respectively, for both normalized fluorescence values. We classified pixels as overlapping when both normalized fluorescence values were above a threshold of 0.3 (inside the gray boxed region). Curves are the average fluorescence between 4 radii of the same colony. **F)** Example of imperfectly symmetrical internal blue rings from a different colony at 52 hours post-plating (see Fig. 7).

**Supplemental Figure 6: Expanding QS colonies in EZ rich defined medium.**

**A)** Colonies obtained from plates with different aTc concentrations, over time. At 100 ng/ml aTc, colonies remained all yellow or with blue center or internal ring fragments. **B)** We used the measurement of color overlap to quantify the spatial segregation of states per colony. We classified pixels as overlapping when both normalized fluorescence values were above a threshold of 0.3. Measurement of overlap was normalized by the colony radius (total number of pixels). Data is from 2 independent experiments. **C)** Fluorescence intensity cross-sectionals of colonies shown in (A). Curves are the average fluorescence between 4 radii of the same colony (see Fig. 7).

**Supplement Figure 7: Schematics of the 3D domain and the discretization of the colony expansion used in numerical simulations. A)** We assume a cone shaped growing colony sits on top of the cylindrical agar pad. The red curves outline a 2D slice from the 3D domain, in the radial (*r*) and height (*z*) direction. Assuming radial symmetry, which is consistent with experimental observations, we used this 2D slice as the domain for the model. The solid line represents the outer boundary (*∂Ω*) while the dashed line represents the inner boundary (*∂Ω*^in^). **B)** Growth is modeled by updating the colony’s shape at even increments in time, *Δt*. At the end of each subinterval, the domain of the colony, *Ω*_1_, is increased by adding a rectangle of height *Δz* and of width equal to that of the colony, and an adjoining right triangle of height *Δz* with base Δ*r*. Here, we discretize time into intervals (*t*_*k*−1_, *t*_*k*_), with *t*_*k*_ = *k* · Δ*t*. **C)** After every increment of time, a new mesh is generated for the expanded colony. The new nodes at the Top and Rim region are initialized by interpolating the solution of the corresponding region from the stretched old mesh (see Fig. 5).

**Supplement Movie 1: Patterning in the expanding NQS toggle colony**. Once aTc drops below the hysteresis point, cells from the top start to switch to the blue state. Sequestration and degradation of aTc leads to the further spreading of the blue wave. Top: aTc profile at different locations of the colony. Middle: 2D colormap of the LacI and TetR profile at different locations of the colony. Bottom three panels on left: normalized LacI and TetR concentration in the top-down view of the colony. Bottom right: Evolution of the effective promoter strength at the top and rim of the colony.

**Supplement Movie 2: Patterning in the expanding QS toggle colony**. Faster growth and balance of the QS signals at the rim leads to cells switching to the blue state. Bistability in the non-growing top leads to preservation of the yellow/blue states, as new cells are pushed out of the actively growing layer. This leads to a stable switching boundary in the radial direction. Top: aTc profile at different locations of the colony. Middle: 2D colormap of the LacI and TetR profile at different locations of the colony. Bottom three panels on left: normalized LacI and TetR concentration in the top-down view of the colony. Bottom right: Evolution of the effective promoter strength at the top and rim of the colony.

