## Supplemental Figure and Table for "Pattern formation and bistability in a synthetic intercellular genetic toggle"

### Supplemental Figures and Tables

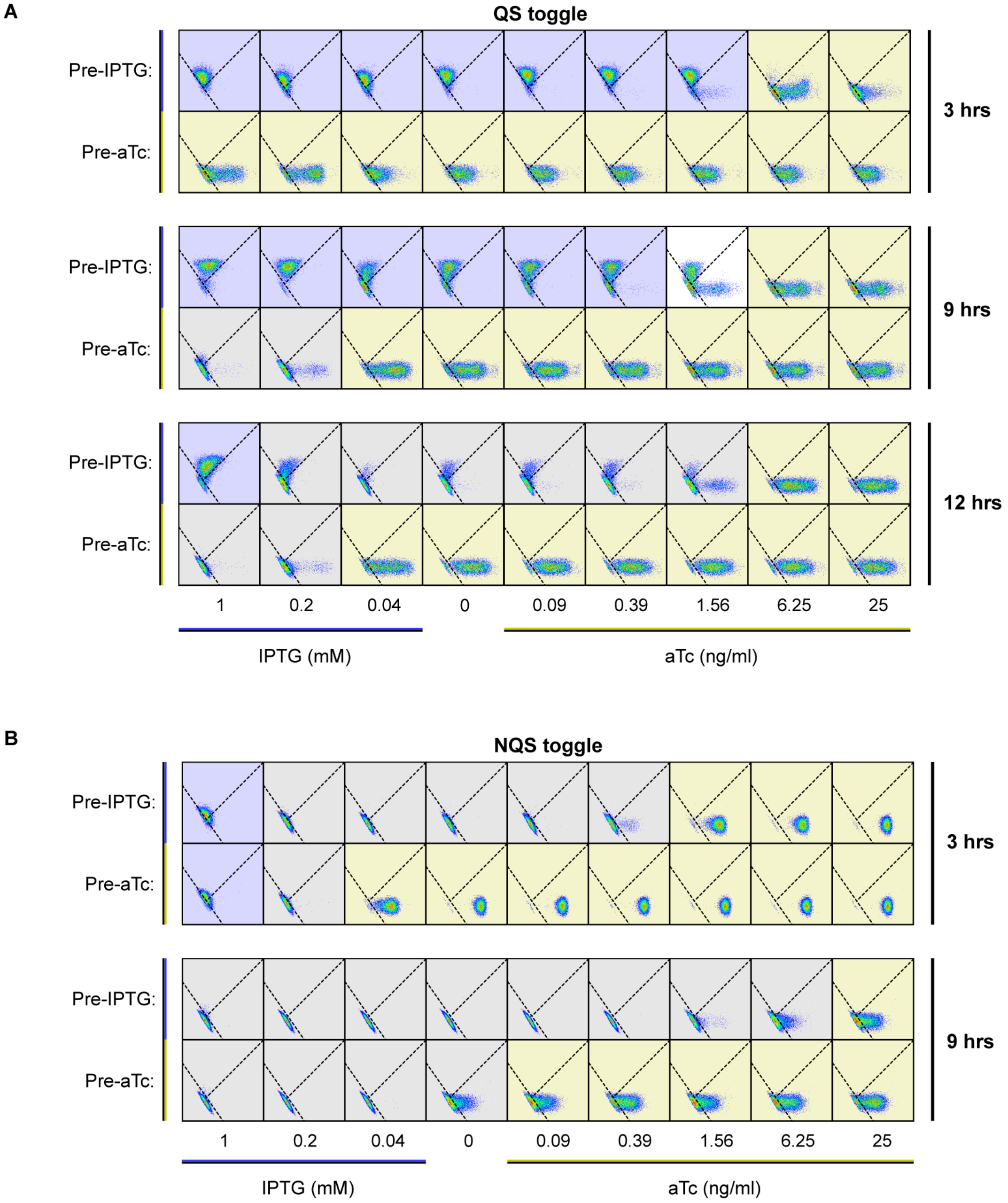


**Supplemental Figure 1: Individual behavior of QS and NQS toggle cells when treated with a single inducer for different duration of times.** Flow cytometry data of QS and NQS toggle cells that were pre-induced with either IPTG or aTc. Each dot is a single cell classified within a gate. Gates were determined with single color and double negative controls. Dashed lines in each plot represent the boundaries between the three distinct gates, which represent cellular states: CFP+ (top gate), YFP+ (bottom-right gate), and OFF (bottom-left gate). Background colors in each plot represent which state the majority of cells are in (>50%): blue color indicates mostly CFP+ cells, yellow plots are mostly YFP+, gray plots are mostly OFF, and white plots indicate cells that are present in multiple states (<50% each). **A)** QS toggle cells pre-treated with IPTG and aTc after growth for 3 (top), 9 (middle), and 12 hours (bottom). **B)** NQS toggle cells pre-treated with IPTG and aTc after growth for 3 (top) and 9 hours (bottom) (see Fig. 2).

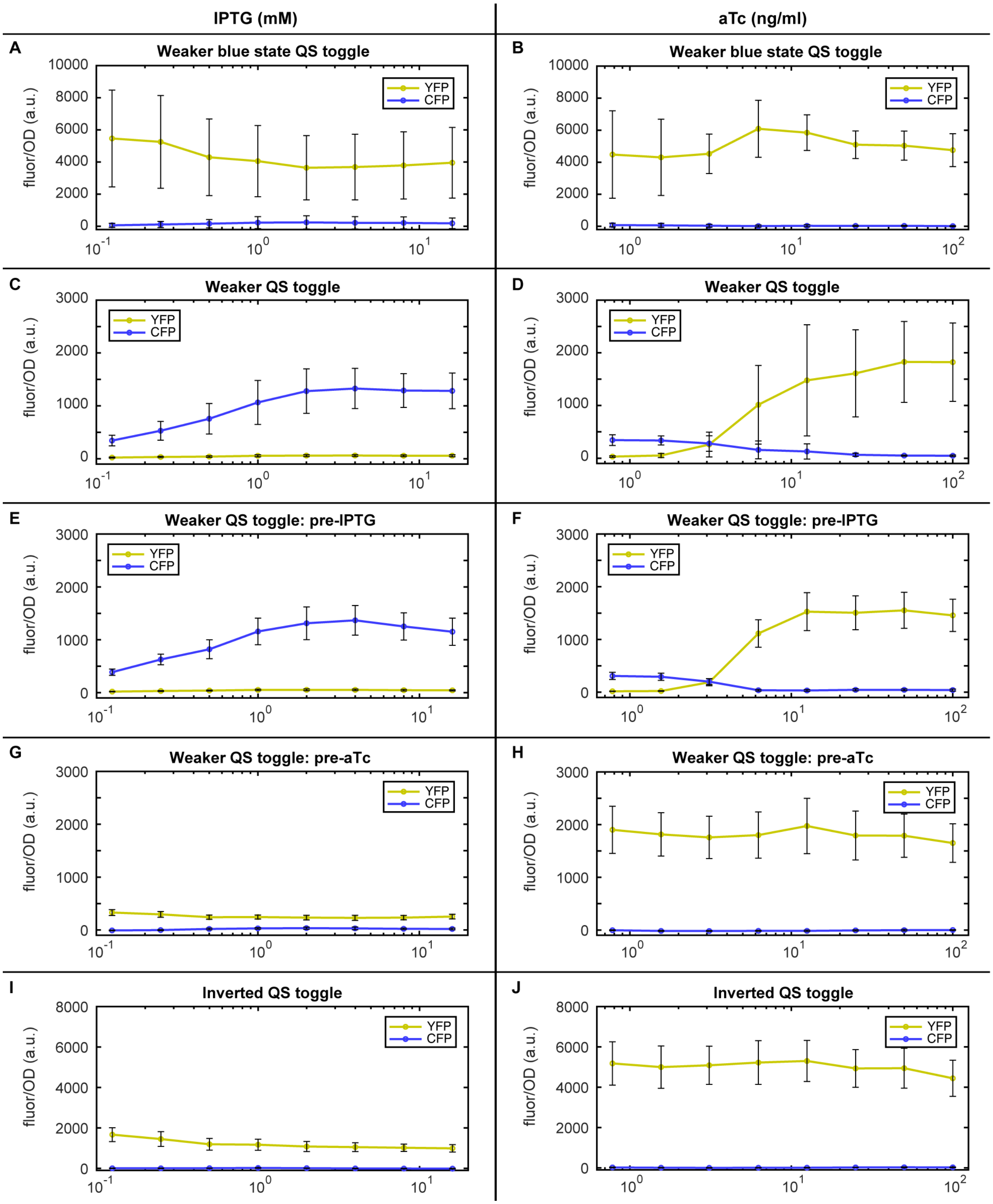

**Supplemental Figure 2: Induction curves of circuit variations of the QS toggle.** We obtained the Weaker Blue State by using lower strength promoters for *tetR* and *cfp* genes. We obtained the Weaker QS toggle by using lower strength promoters for the circuit genes, except both QS genes which were kept the same. We obtained the Inverted QS toggle by using the same lower strength promoters but reversing the QS network connected to each state: now, the rhlR/I network activates the yellow state, and cinR/I the blue state. **A, B) I**nduction curves of Weaker Blue State QS toggle with IPTG (A) and aTc (B) in liquid culture. **C, D)** Induction curves of Weaker QS toggle with IPTG (C) and aTc (D) in liquid culture. **E, F)** Induction curves of Weaker QS toggle with IPTG (E) and aTc (F) in liquid culture, in which cells were pre-treated with IPTG. **G, H)** Induction curves of weaker QS toggle with IPTG (G) and aTc (H) in liquid culture, in which cells were pre-treated with aTc. **I, J)** Induction curves of Inverted QS toggle with IPTG (I) and aTc (J) in liquid culture. Lines represent the average fluorescence and error bars represent the standard deviation of 3 technical replicates for at least 3 independent experiments (see Fig. 1).

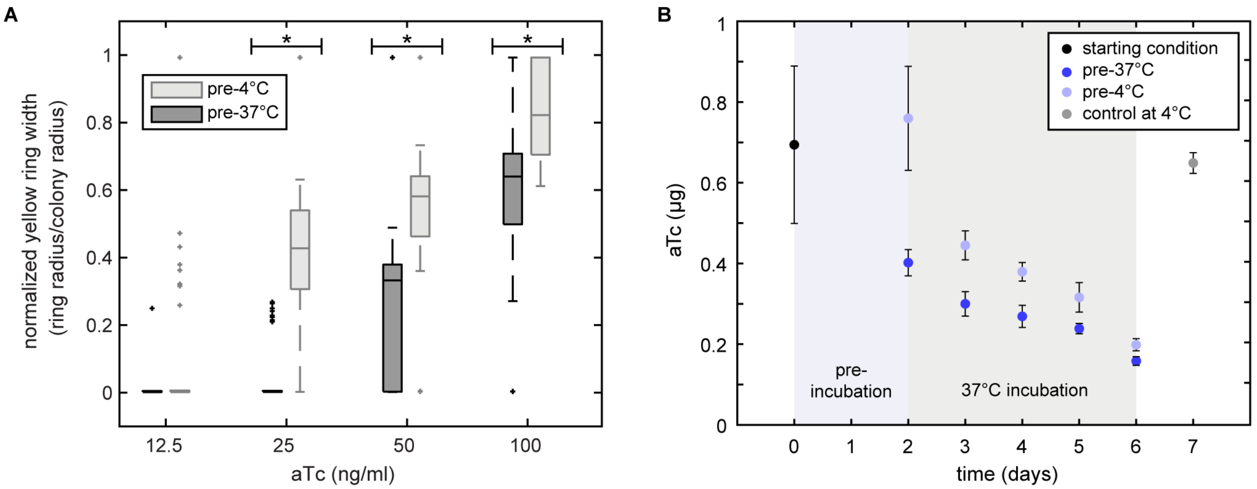


**Supplemental Figure 3: Incubation at 37°C causes aTc degradation. A)** Yellow ring width measured from QS toggle colonies grown in aTc plates pre-incubated either at 4°C or 37°C for 48 hours prior to plating. Then, we plated and grew cells as shown in Fig. 3A. These values represent yellow ring widths from 76h post-plating. The 37°C plates showed a significant decrease in width in comparison to its 4°C counterparts, except at 12.5 ng/ml (*$p < 0.01$, Mann-Whitney non-parametric test). Data is from 5 independent experiments. Pre-4°C colonies are included in Fig. 3B. **B)** ATc quantification from LB agar extracts (in the absence of cells) with an HPLC. At time 0, aTc was quantified before any incubation. Samples were divided in two groups: pre-incubation for 48 hours at 37°C (bright blue), or 4°C (light blue) to recreate the experimental timeline in (A). On day 2, we incubated both groups at 37°C until day 6 to also recapitulate the experimental setup. As a control (gray), LB agar + aTc samples were kept at 4°C throughout the entire test and measured on day 7. Data represents mean ± standard deviation of 3 independent experiments (see Fig. 3).

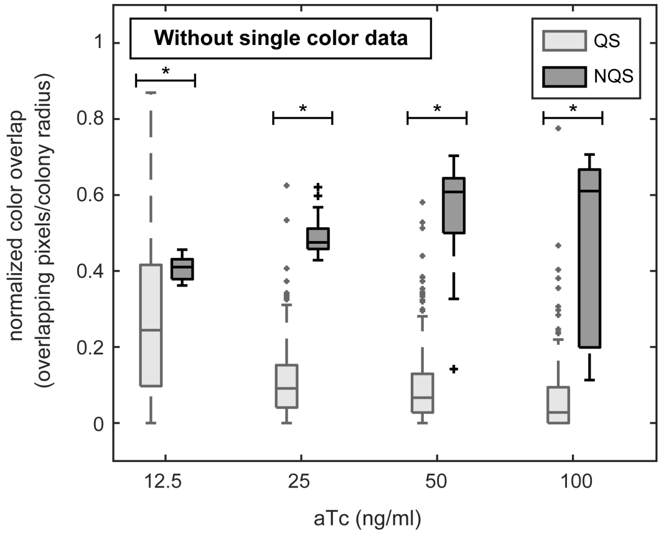


**Supplemental Figure 4: Quantification of pixel overlap for QS toggle (light gray) and NQS toggle (dark gray) colonies without single color data.** We selected only colonies that have both colors present for at least 25% of the radius. We normalized the number of overlapping pixels by the colony radius (total pixels) ($p < 0.01$, Mann-Whitney non-parametric test). Data for each QS toggle test contains over 138 colonies from at least 11 independent experiments, while data for each NQS test contains at least 5 colonies from 2 independent experiments (see Fig. 4).

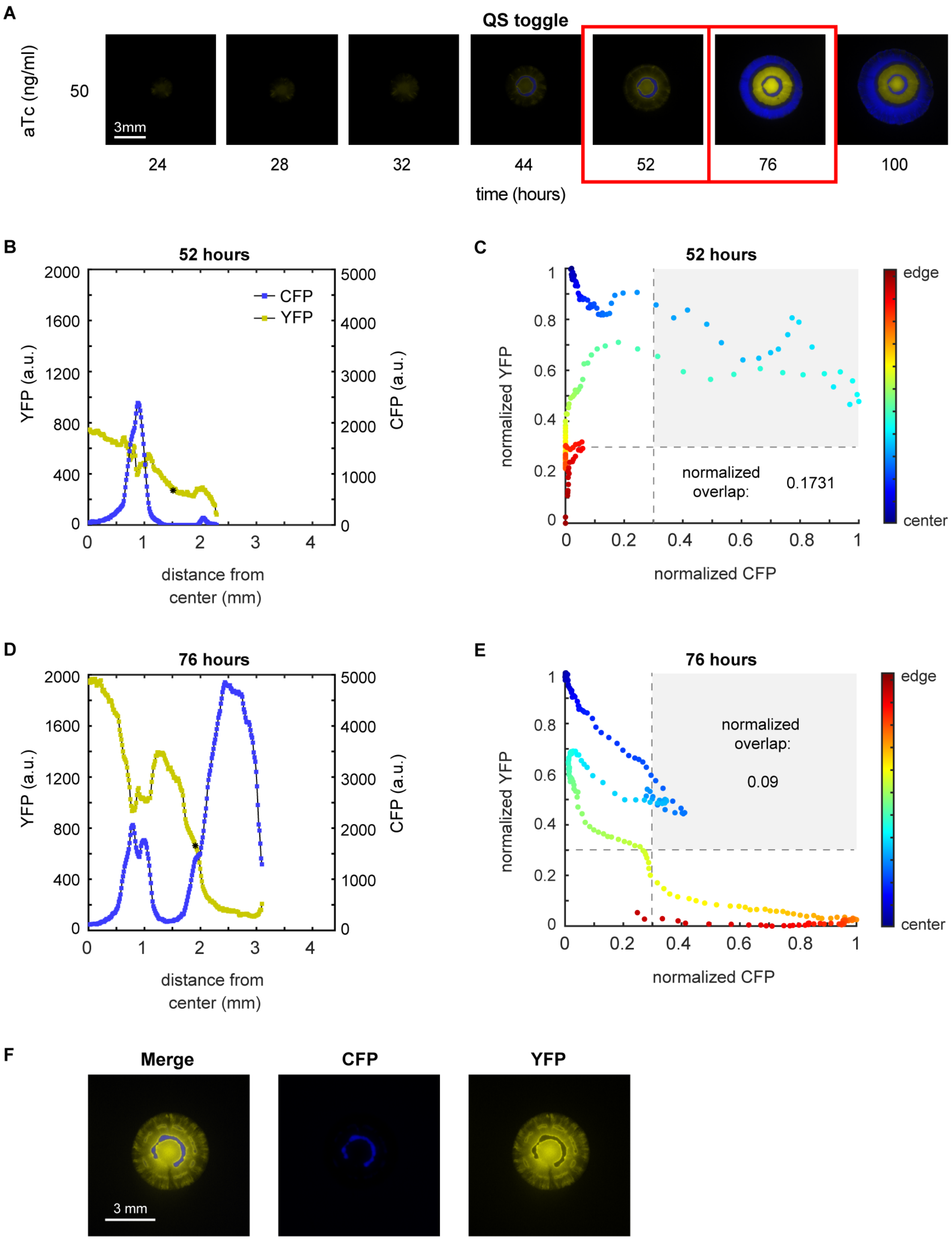


**Supplemental Figure 5: Multiple blue rings are also observed in LB agar QS colonies. A)** Example of colony obtained from a 50 ng/ml aTc plate, over time. **B, C)** Fluorescence intensity cross-section at 52h and 76h, respectively, shown in (A). Curves are the average fluorescence between 4 radii of the same colony. **D, E)** We plotted each colony pixel from the images at 52h and 76h, respectively, for both normalized fluorescence values. We classified pixels as overlapping when both normalized fluorescence values were above a threshold of 0.3 (inside the gray boxed region). Curves are the average fluorescence between 4 radii of the same colony. **F)** Example of imperfectly symmetrical internal blue rings from a different colony at 52 hours post-plating (see Fig. 7).

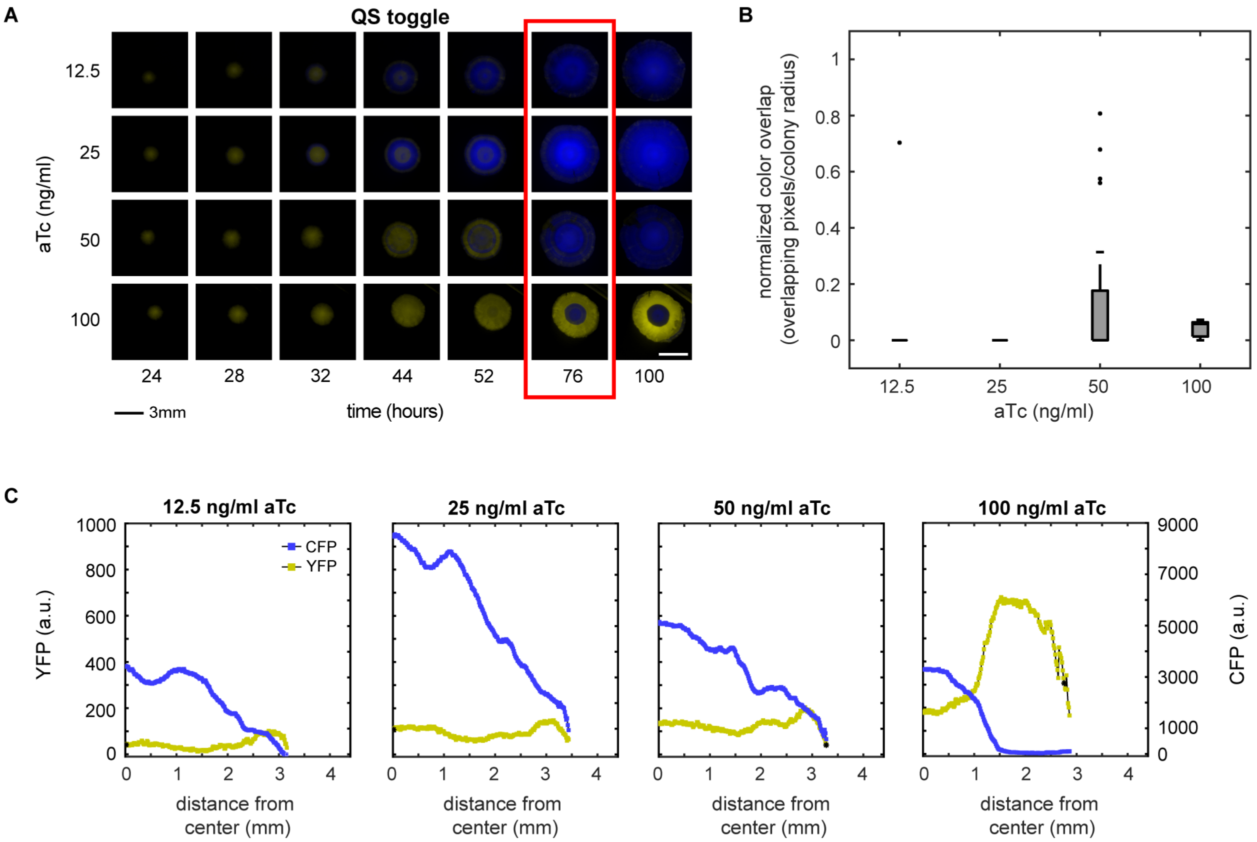


**Supplemental Figure 6: Expanding QS colonies in EZ rich defined medium. A)** Colonies obtained from plates with different aTc concentrations, over time. At 100 ng/ml aTc, colonies remained all yellow or with blue center or internal ring fragments. **B)** We used the measurement of color overlap to quantify the spatial segregation of states per colony. We classified pixels as overlapping when both normalized fluorescence values were above a threshold of 0.3. Measurement of overlap was normalized by the colony radius (total number of pixels). Data is from 2 independent experiments. **C)** Fluorescence intensity cross-sectionals of colonies shown in (A). Curves are the average fluorescence between 4 radii of the same colony (see Fig. 7).

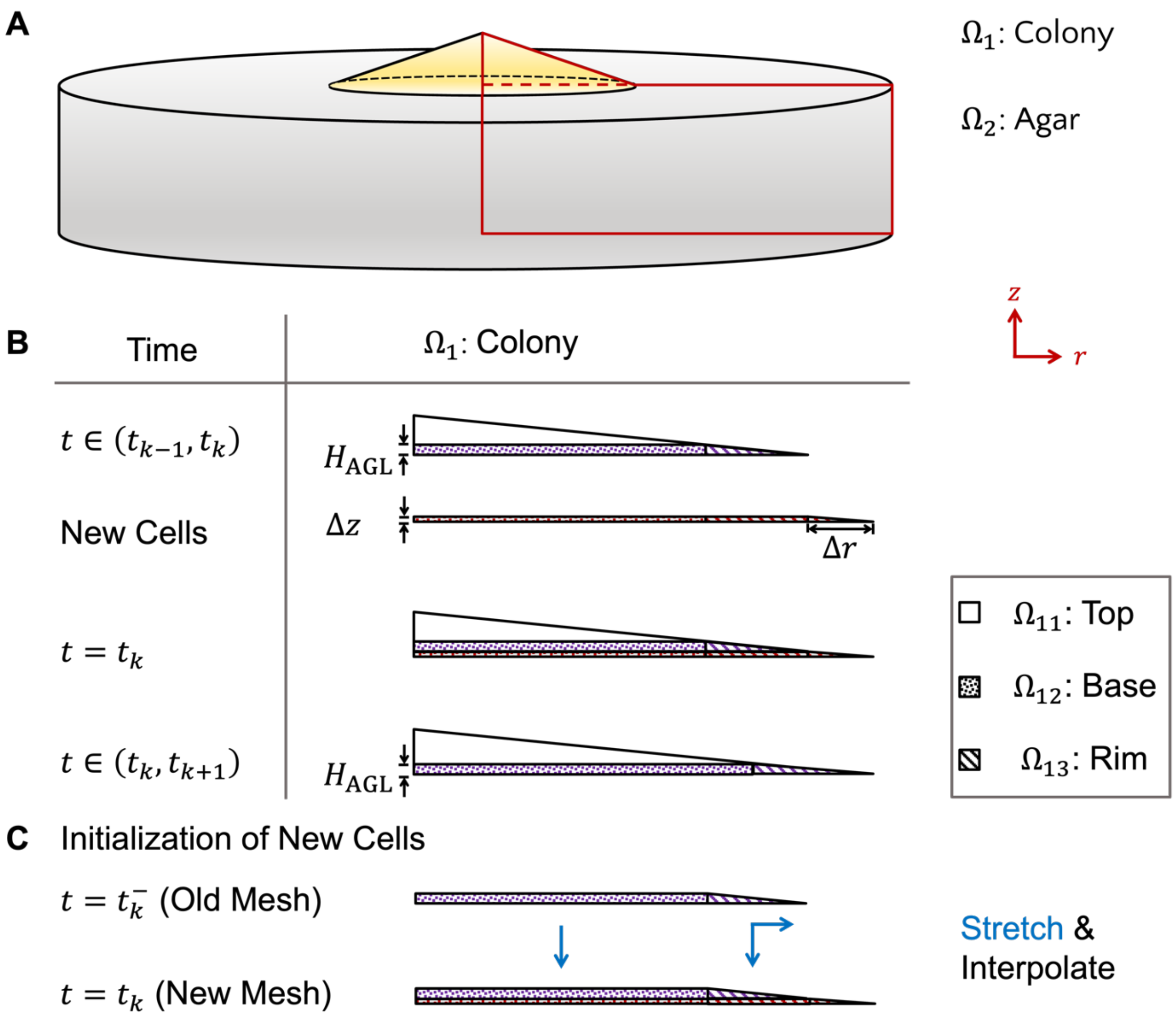


**Supplement Figure 7: Schematics of the 3D domain and the discretization of the colony expansion used in numerical simulations. A)** We assume a cone shaped growing colony sits on top of the cylindrical agar pad. The red curves outline a 2D slice from the 3D domain, in the radial ($r$) and height ($z$) direction. Assuming radial symmetry, which is consistent with experimental observations, we used this 2D slice as the domain for the model. The solid line represents the outer boundary ($\partial\Omega$) while the dashed line represents the inner boundary ($\partial\Omega^{\mathrm{in}}$). **B)** Growth is modeled by updating the colony's shape at even increments in time, $\Delta t$. At the end of each subinterval, the domain of the colony, $\Omega_1$, is increased by adding a rectangle of height $\Delta z$ and of width equal to that of the colony, and an adjoining right triangle of height $\Delta z$ with base $\Delta r$. Here, we discretize time into intervals $(t_{k-1}, t_k)$, with $t_k = k \cdot \Delta t$. **C)** After every increment of time, a new mesh is generated for the expanded colony. The new nodes at the Top and Rim region are initialized by interpolating the solution of the corresponding region from the stretched old mesh (see Fig. 5).

**Supplement Table 1: List of parameters used in simulations unless otherwise mentioned in the text.**

| Type | Parameters | Description | Value | Units |
|---|---|---|---|---|
| NQS | $a_1$ | LacI max. production rate | $100$ | $\mathrm{nM} \cdot \mathrm{min}^{-1}$ |
| | $a_2$ | TetR max. production rate | $250$ | $\mathrm{nM} \cdot \mathrm{min}^{-1}$ |
| | $u_0$ | aTc Concentration (low/medium/high) | $15/30/60$ | $\mathrm{nM}$ |
| | $k_+$ | aTc-TetR binding | $0.06$ | $\mathrm{nM}^{-1} \cdot \mathrm{min}^{-1}$ |
| | $k_-$ | aTc-TetR unbinding | $0.006$ | $\mathrm{min}^{-1}$ |
| QS | $a_1$ | LacI max. production rate | $110$ | $\mathrm{nM} \cdot \mathrm{min}^{-1}$ |
| | $a_2$ | TetR max. production rate | $210$ | $\mathrm{nM} \cdot \mathrm{min}^{-1}$ |
| | $a_3$ | Effective C14 max. production rate | $250$ | $\mathrm{nM} \cdot \mathrm{min}^{-1}$ |
| | $a_4$ | Effective C4 max. production rate | $2000$ | $\mathrm{nM} \cdot \mathrm{min}^{-1}$ |
| | $u_0$ | aTc Concentration (low/medium/high) | $24/30/36$ | $\mathrm{nM}$ |
| | $k_+$ | aTc-TetR binding | $0.006$ | $\mathrm{nM}^{-1} \cdot \mathrm{min}^{-1}$ |
| | $k_-$ | aTc-TetR unbinding | $0.0006$ | $\mathrm{min}^{-1}$ |
| NQS/QS | $\gamma_1$ | LacI degradation rate | $\ln 2/7$ | $\mathrm{min}^{-1}$ |
| | $\gamma_2$ | TetR degradation rate | $\ln 2/7$ | $\mathrm{min}^{-1}$ |
| | $\gamma_3$ | C14 degradation rate | $\ln 2/24$ | $\mathrm{h}^{-1}$ |
| | $\gamma_4$ | C4 degradation rate | $\ln 2/24$ | $\mathrm{h}^{-1}$ |
| | $\gamma_5$ | aTc degradation rate | $\ln 2/48$ | $\mathrm{min}^{-1}$ |
| | $\theta_x$ | IC50 of LacI for $\mathrm{P_{Lac}}$ | $500$ | $\mathrm{nM}$ |
| | $\theta_y$ | IC50 of TetR for $\mathrm{P_{Tet}}$ | $500$ | $\mathrm{nM}$ |
| | $\theta_g$ | EC50 of C14 for $\mathrm{P_{Cin/Tet}}$ | $100$ | $\mathrm{nM}$ |
| | $\theta_h$ | EC50 of C4 for $\mathrm{P_{Rhl/Lac}}$ | $100$ | $\mathrm{nM}$ |
| | $D_1^a$ | aTc diffusion in agar | $400$ | $\mathrm{\mu m}^2 \cdot s^{-1}$ |
| | $D_1^c$ | aTc diffusion in colony | $40$ | $\mathrm{\mu m}^2 \cdot s^{-1}$ |
| | $D_3^a$ | C14 diffusion in agar | $10^2$ | $\mathrm{\mu m}^2 \cdot s^{-1}$ |
| | $D_3^c$ | C14 diffusion in colony | $10$ | $\mathrm{\mu m}^2 \cdot s^{-1}$ |
| | $D_4^a$ | C4 diffusion in agar | $10^3$ | $\mathrm{\mu m}^2 \cdot s^{-1}$ |
| | $D_4^c$ | C4 diffusion in colony | $10^2$ | $\mathrm{\mu m}^2 \cdot s^{-1}$ |
| | $\kappa$ | Colony Aspect Ratio | $1:10$ | - |
| | $v_r$ | Radial growth rate | $40$ | $\mathrm{\mu m/h}$ |
| | $v_h$ | Vertical growth rate | $4$ | $\mathrm{\mu m/h}$ |
| | $H_{\mathrm{AGL}}$ | Thickness of AGL | $10$ | $\mathrm{\mu m}$ |
| | $T$ | Time takes to grow $H_{\mathrm{AGL}}$ | $2.5$ | $\mathrm{h}$ |
| | $\gamma_d$ | dilution from linear growth | $0.4$ | $\mathrm{h}^{-1}$ |
